# Microenvironmental Fingerprint of Maturing Tau Condensates from Spectral and Lifetime Phasor Analysis of a Solvatochromic Dye

**DOI:** 10.64898/2026.11.07.765020

**Authors:** Leandro Cruz Rodríguez, Yarne Vankerkhoven, Sonny Brebels, Ilse Dewachter, Enrico Carlon, Jelle Hendrix

## Abstract

Solvatochromic dyes respond to changes in their local environment through variations in both spectral properties and fluorescence lifetime. These responses arise from the probe’s excited-state behavior, but the sensitivity of each response to a given environment can differ. In this work, we developed a multimodal phasor-based frame-work for two-photon spectral and fluorescence lifetime imaging analysis. To validate the framework, we imaged the solvatochromic dye ACDAN in water, glycerol, and ethanol. ACDAN showed clear spectral differences among the three solvents, whereas its fluorescence lifetime separated water from both organic solvents but not glycerol from ethanol, indicating that the two observables respond differently to the same environmental change. We then applied ACDAN to a well-described model system, the maturation of liquid–liquid phase-separated condensates of the microtubule-binding, neurodegeneration-related protein Tau into later-stage assemblies. Across Tau samples incubated for 1, 4, and 48 h, a progressive decrease in the ACDAN spectral phasor angle *θ* and increase in its phase-derived lifetime *τ_P_* suggested increasingly restricted dipolar relaxation in the probe environment. Differences among condensates were apparent by 4 h, whereas pronounced spatial heterogeneity within assemblies was observed at 48 h. Lastly, we evaluated the multimodal framework on Tau assemblies labeled with the fluorogenic probe Thioflavin T. As expected, this dye predominantly sensed its environment through its fluorescence lifetime, with comparatively limited spectral changes, whereas ACDAN reported microenvironmental changes in both coordinates. Together, this framework combines spectral and lifetime descriptors to characterize changes in the fluorescence response across Tau assembly maturation while retaining their spatial distribution.

## MAIN

Solvatochromic dyes are used in fluorescence microscopy because their fluorescence characteristics respond to the physicochemical properties of their immediate environment. These properties include local polarity, hydration, molecular confinement, and solvent reorganization around the excited fluorophore, and they can affect the absorption and emission spectra, fluorescence quantum yield, and fluorescence lifetime of the dye.^1–4^ The DAN (6-acyl-2-dimethylaminonaphthalene) family is among the most extensively used classes of solvatochromic dyes. Following excitation, solvent molecules reorganize around the increased dipole moment of the probe, and the extent and kinetics of this relaxation subsequently influence its fluorescence emission.^1,2,5–7^ LAURDAN (6-lauroyl-2-dimethylaminonaphthalene) is widely used to report lipid membrane properties^2^ and ACDAN (6-acetyl-2-dimethylaminonaphthalene, Supplementary Fig. S1a) is a water-soluble variant that senses changes in water organization, molecular crowding, and confinement in solutions, cells, and tissues. ^3,8–10^

The phasor approach to fluorescence provides a graphical framework for analyzing spectral and lifetime information from fluorescence microscopy data.^11–13^ Its theoretical basis can be traced to earlier studies of phase- and modulation-fluorometry in heterogeneous fluorescence systems, which established the use of phase, modulation, and Cartesian co-ordinates *G* and *S* to represent fluorescence decays.^14,15^ Related graphical approaches were subsequently developed for single-frequency fluorescence lifetime imaging microscopy and frequency-domain lifetime analysis.^16,17^ In phasor-based FLIM, the fluorescence decay associated with each pixel is represented by Cartesian coordinates (*G, S*) at a selected modulation frequency or Fourier harmonic. Monoexponential decays lie on the universal semicircle, whereas multicomponent or heterogeneous decays generally fall within its interior. ^14,15,18^ The reciprocal relationship between phasor space and image space allows selected phasor populations to be mapped back onto the corresponding pixels in the sample image.^12,18^ Spectral phasor analysis, on the other hand, applies an analogous Fourier transformation to the emission spectrum recorded at each pixel. ^19^ In this representation, the phasor angle primarily reports the position of the emission band within the acquisition window, whereas the modulus provides information about its spectral width.^13,19^ As in phasor-FLIM, selected spectral phasor populations can be mapped back to the pixels from which they were calculated. Neither transformation requires a model of the spectrum or of the decay. The strategy has also been extended to low-signal and single-particle measurements. For example, particle-based phasor-FLIM-FRET has been used to resolve protein–protein interactions inside single viral particles.^20^

Combined spectral and lifetime analysis can be used to characterize complex biological samples.^21^ Malacrida and colleagues combined spectral and lifetime phasor measurements of LAURDAN from the same cellular plane to distinguish membrane-polarity effects from dipolar relaxation.^21^ Scipioni et al. developed spectrally resolved FLIM with multidimensional phasor analysis to disentangle overlapping spectral and lifetime signatures.^22^ Haas et al. applied multidimensional phasor demixing to distinguish three FRET biosensors expressed in the same living cells, allowing their responses to be monitored in parallel.^23^

Biomolecular condensates provide a relevant setting for combined spectral and lifetime analysis. In these systems, macromolecules become concentrated within a dense liquid phase that coexists with a surrounding dilute phase.^24^ ACDAN-based spectral and lifetime measurements have detected differences between these phases and changes associated with protein concentration, solution conditions, and condensate material properties. ^25–29^ The fluorescence response of ACDAN can therefore be used to follow changes in the local environment of condensates without requiring the probe to recognize a specific protein structure. Tau provides a useful system for investigating these environments during condensate maturation. The protein can undergo liquid–liquid phase separation (LLPS) under several *in vitro* conditions, and the resulting condensates can change in molecular organization and dynamics during incubation.^30,31^ Prolonged residence in the condensed phase has been associated with oligomerization and the appearance of disease-related Tau conformations in some experimental systems.^30^ Under aggregation-promoting conditions, particularly in the presence of polyanionic cofactors such as heparin, phase separation can also accelerate the formation of fibrillar assemblies. ^32^ Recent FLIM measurements of an engineered Tau variant also revealed differences between condensate interfaces and interiors during maturation.^33^ A recent review of DAN probes in condensates proposed the joint analysis of spectral and lifetime information at the pixel level as a way to separate contributions from polarity, hydration, and intermolecular interactions. ^34^ Spectral and lifetime phasor analysis of environment-sensitive probes has been used to follow the maturation of *β*-lactoglobulin condensates formed under polymer crowding, where aging was accompanied by reduced molecular mobility and the emergence of spatial heterogeneity within individual condensates. ^35^ The evolution of the local environment within and between individual Tau assemblies during maturation is less well described.

In this work, we combined spectral imaging and FLIM in a phasor-based framework to investigate microenvironmental changes during Tau condensate maturation using ACDAN. We first characterized the spectral and lifetime responses of ACDAN in reference solvents. The solvatochromism of ACDAN and related DAN probes is well established, ^1,3,8^ and these measurements were used to validate the framework on a system with a known solvent dependence before applying it to protein assemblies. We then applied the same framework to Tau condensates and later-stage assemblies obtained at different incubation times under aggregation-promoting conditions. Finally, we analyzed Thioflavin T using the same workflow to compare the spectral and lifetime responses of the two probes.

## Results

### Spectral and lifetime phasor analysis of ACDAN in water

Since we aimed to use two-photon excitation-based fluorescence imaging of ACDAN, we first characterized the dye in detail by separate spectral and lifetime imaging analysis in water (Fig. 1).

**Figure 1:**
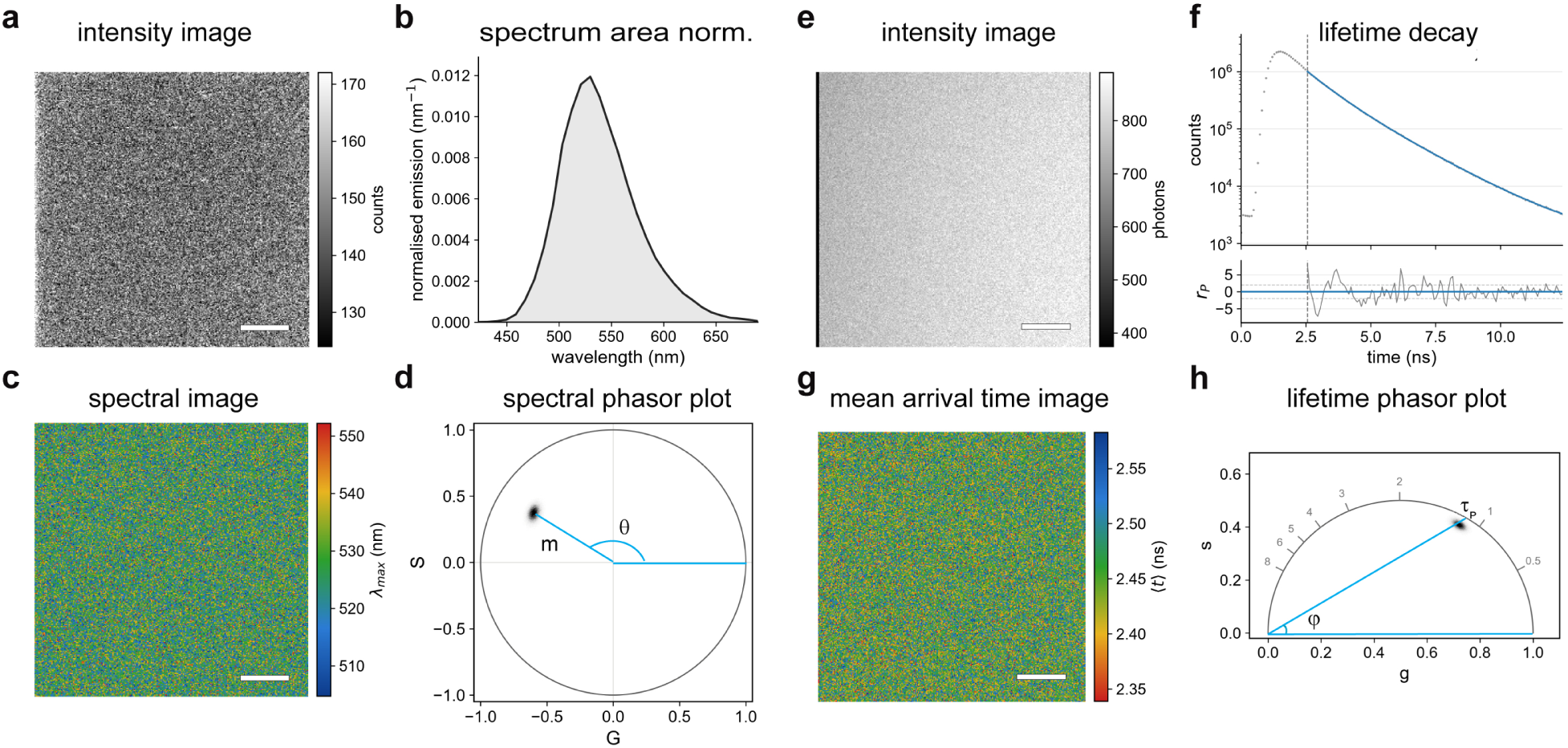
Spectral and lifetime phasor analysis of ACDAN in water. The left block (a–d) shows the hyperspectral acquisition and the right block (e–h) the FLIM acquisition of the same field of view, each read row by row. (a) Intensity image, summed over all spectral channels. (b) Pixel-averaged emission spectrum. (c) Spatial map of the pixel-wise emission maximum, *λ*_max_. (d) Spectral phasor of all valid pixels. The vector from the origin to the cluster centroid defines the phasor angle *θ* and the modulus *M*_sp_; only *θ* is used in the joint analysis. (e) FLIM intensity image of the same field of view. (f) Spatially integrated fluorescence decay with the fitted two-component model and weighted residuals *r_P_*; the dashed line marks the start of the fitting window. (g) Image of the mean photon arrival time. This quantity includes the instrument response offset and is not a lifetime. (h) Lifetime phasor of all valid pixels relative to the universal semicircle. The phase angle *φ* defines *τ_P_*, read where the radial direction meets the semicircle, and the modulus defines *τ_M_*; numbers along the semicircle indicate lifetime in nanoseconds at the repetition rate used. Scale bars: 5 µm.

Summed over all spectral channels and image frames, the intensity image of ACDAN in water was spatially uniform (Fig. 1a), with an average of 147 counts per pixel distributed over 31 emission channels spanning 423–690 nm. The pixel-averaged emission spectrum peaked at 525.1 ± 0.4 nm and had a full width at half maximum of 74.8 nm (Fig. 1b), consistent with previous measurements of ACDAN in aqueous solution. ^8,36^ The pixel-wise emission maximum is shown in Fig. 1c, with a median of 527.4 nm and an interquartile range of 11.4 nm.

The spectral phasor transformation provides a different description of the same data by representing each pixel through its first-harmonic coordinates (*G*_sp_*, S*_sp_). ACDAN in water formed a compact population within the spectral phasor plot (Fig. 1d). The phase angle *θ* = atan2(*S*_sp_*, G*_sp_) mod 2*π* describes the position of the emission distribution within the spectral window, whereas the modulus 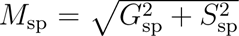 contains information related to spectral width.^19^ For ACDAN in water, *θ* = 2.585 rad with an interquartile range of 0.077 rad.

Next, we performed FLIM of the same sample, which provided the temporal response of ACDAN, with an average of 798 photons per pixel (Fig. 1e). The spatially integrated fluorescence decay was better described by a biexponential than by a monoexponential model after accounting for the additional fit parameters (ΔBIC *>* 10^4^), yielding lifetime components of 0.90 and 1.81 ns, although some structure remained in the residuals (Fig. 1f). The mean photon arrival time of each pixel is shown in Fig. 1g.

The same decay data were then also analyzed in the phasor domain. Each pixel was represented by its calibrated (*G*_lf_ *, S*_lf_) coordinates relative to the universal semicircle (Fig. 1h). The phase angle *φ* yielded a phase lifetime of *τ_P_* = 1.12 ns, while the phasor modulus yielded a modulation lifetime of *τ_M_* = 1.34 ns. The pixel population was displaced toward the interior of the universal semicircle, with *κ* = *τ_M_ /τ_P_* = 1.20. This behavior is consistent with a non-monoexponential fluorescence decay and agrees with the temporal heterogeneity indicated independently by the decay fitting.^11,12,18^

These measurements define the two descriptors used in the subsequent multimodal analysis. The spectral coordinate *θ* is the phase angle of the first-harmonic spectral phasor, whereas *τ_P_* is the phase lifetime calculated from the phase angle of the first-harmonic life-time phasor. The spectral modulus and the modulation lifetime are not retained. In what follows, we use the resulting (*θ, τ_P_*) plane as a derived feature space for comparing fluorescence responses.^12,18,19^

### Photophysical response of ACDAN in reference solvents

To investigate the dependence of ACDAN’s spectroscopic properties on its physicochemical environment, we next characterized the dye in water, ethylene glycol, glycerol, and ethanol, which differ in polarity, hydrogen-bonding capacity, and viscosity. ^1,3,8^ All four solvents were measured in cuvettes, and water, glycerol, and ethanol were also imaged.

Cuvette-based measurements showed that the fluorescence quantum yield of ACDAN was 0.20, 0.67, 0.77, and 0.84 in water, ethylene glycol, glycerol, and ethanol, respectively, with corresponding fluorescence lifetimes of 1.24, 2.75, 3.26, and 3.36 ns (Supplementary Tables S1 and S2). The apparent non-radiative rate varied more strongly between these solvents, from 0.048 ns*^−^*^1^ in ethanol to 0.64 ns*^−^*^1^ in water, than the apparent radiative rate, which ranged from 0.16 to 0.25 ns*^−^*^1^. Glycerol and ethanol showed similar fluorescence lifetimes despite their markedly different viscosities, while their emission spectra remained distinct. Conversely, ethylene glycol and glycerol had similar emission maxima but different fluorescence lifetimes (Supplementary Table S2). These comparisons show that spectral shifts and excited-state deactivation do not respond identically to changes in solvent environment. Using the same experimental conditions and microscope settings as in Fig. 1, we then performed spectral and lifetime imaging (Fig. 2a, top row). Consistent with its lower fluorescence quantum yield, ACDAN in water produced substantially lower fluorescence intensities than in either organic solvent. Differences in photon number are also relevant to the phasor distributions, since lower photon counts increase pixel-wise dispersion and must be accounted for when comparing population widths.

**Figure 2:**
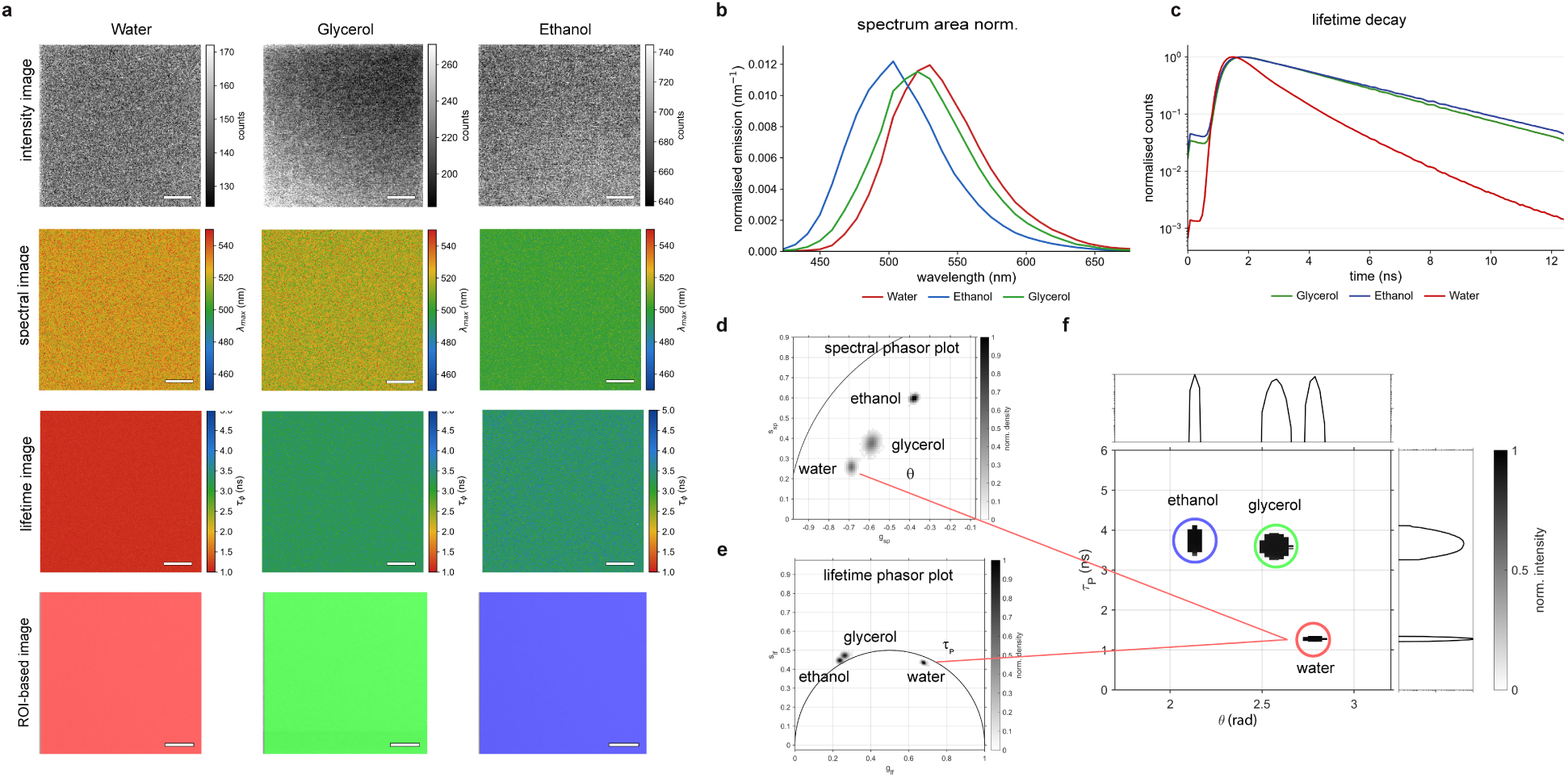
ACDAN exhibits a different spectral and lifetime signature in different solvents. (a) ACDAN in water, glycerol and ethanol imaged under identical concentrations and imaging settings. Rows, top to bottom: total intensity after spectral imaging; pixel-wise emission maximum, *λ*_max_; pixel-wise phase lifetime, *τ_P_*; pixel-wise assignment to the regions of interest defined in (f). The second and third rows use a color scale shared across solvents. (b) Averaged emission spectra, area normalized. (c) Average fluorescence decays, normalized. (d) Spectral phasor and (e) lifetime phasor of all valid pixels. The three solvents are resolved in (d), whereas glycerol and ethanol overlap in (e). (f) Intensity-weighted density in the joint (*θ, τ_P_*) plane. The three regions of interest (ROIs) are defined once on this distribution and propagated pixel-wise to the image space in (a) without re-thresholding. Scale bars: 5 µm. propagated back to the corresponding image pixels (Fig. 2a, bottom row), linking the joint spectral and lifetime response to its spatial origin.

The emission spectrum showed a clear solvent dependence, shifting from approximately 495 nm in ethanol to 510 nm in glycerol and 525 nm in water (Fig. 2b). The corresponding pixel-wise spectral maps were likewise different in the three media (Fig. 2a, second row), and their populations clearly occupied distinct regions in the spectral phasor plot (Fig. 2d).

The fluorescence lifetime ordered the same solvents differently. The decay of ACDAN in water was clearly separated from those in the two organic solvents, which were almost indistinguishable from each other (Fig. 2c), as confirmed by the lifetime phasor (Fig. 2e) and corresponding *τ_P_* maps (Fig. 2a, third row).

In the combined (*θ, τ_P_*) plane, the three solvents occupied distinct regions (Fig. 2f). Glycerol and ethanol were distinguished mainly along the spectral coordinate, whereas water also differed markedly in fluorescence lifetime. Three ROIs were defined in this plane and propagated back to the corresponding image pixels (Fig. 2a, bottom row), linking the joint spectral and lifetime response to its spatial origin.

### Multimodal mapping of early-stage Tau condensates

Next, we applied the same analysis to Tau condensates prepared with PEG as a crowding agent and heparin as a polyanionic cofactor that promotes Tau fibrillation. ^32^ Within 1 h of incubation at 37 °C, the intensity image showed a rounded condensate with higher ACDAN fluorescence than the surrounding phase and several bright punctate features, particularly near its periphery (Fig. 3a). The spectral map showed lower *θ* within the condensate than in the surrounding phase, with several localized foci displaying a further decrease. The *τ_P_* map showed longer lifetimes within the condensate and a region of higher *τ_P_* along its lower-right boundary. The small foci were less distinct in the lifetime map. However, because the two modalities were acquired sequentially over different durations, their different appearance may partly reflect motion during acquisition.

**Figure 3:**
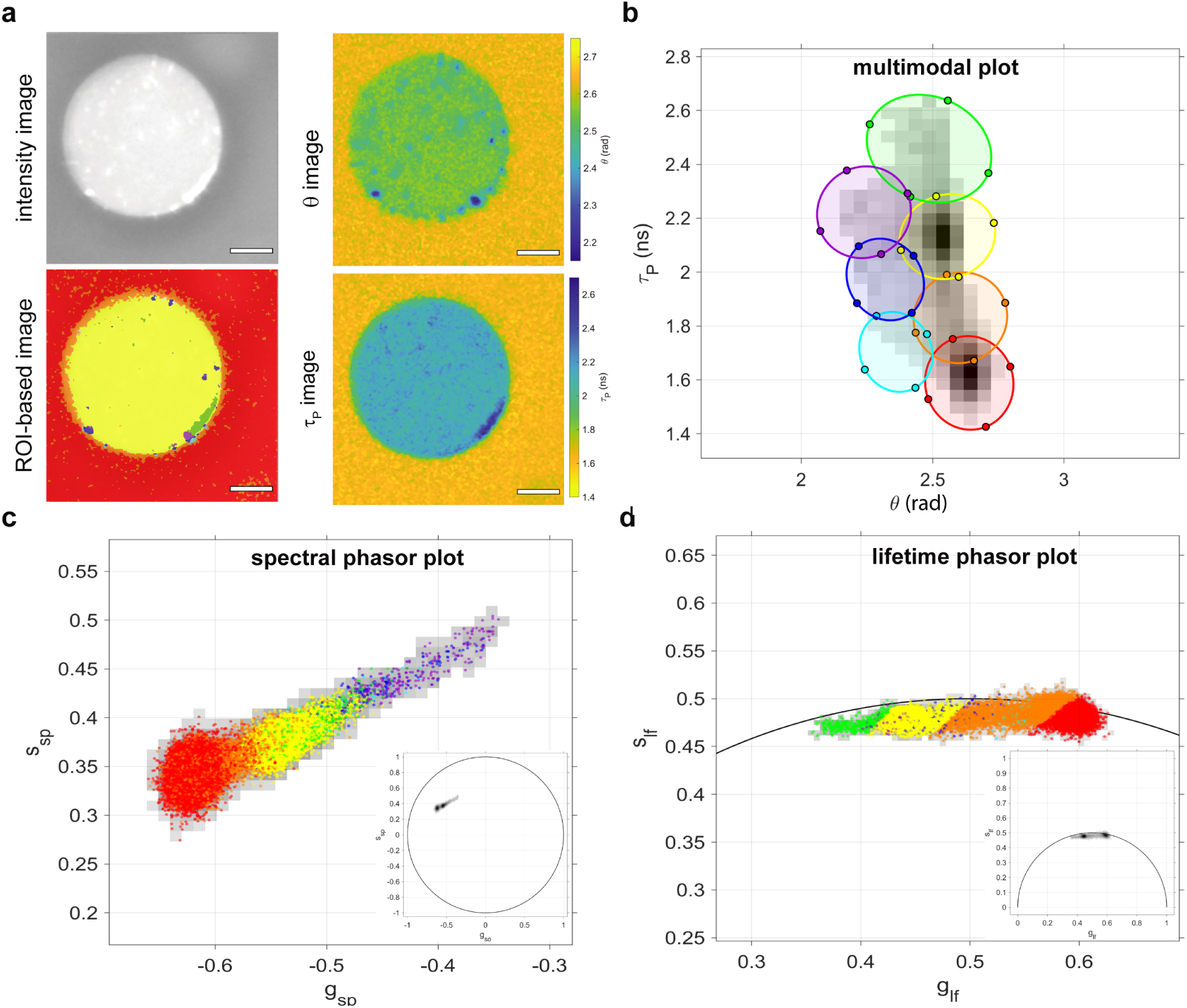
Multimodal mapping of a Tau condensate after 1 h of incubation. (a) The same field of view rendered in four ways. Top left: hyperspectral intensity, summed over all emission channels. Top right: spectral phasor angle *θ*, pixel-wise. Bottom right: phase lifetime *τ_P_*, pixel-wise. Bottom left: pixel-wise assignment to the regions of interest defined in (b). The *θ* and *τ_P_* maps are displayed over the ranges spanned by the corresponding axes in (b). (b) Intensity-weighted density of all valid pixels in the combined (*θ, τ_P_*) plane. Seven elliptical regions of interest are defined on this distribution and propagated pixel-wise to image space in (a) without re-thresholding. Filled circles mark the semi-axis endpoints of each ellipse. (c) Spectral phasor and (d) lifetime phasor of the same pixels, colored by ROI assignment. Gray shading is the density of all valid pixels, and the black curve in (d) is the universal semicircle. Insets show the full phasor space, with the region displayed in the main panel indicated. Scale bars: 5 µm.

The combined (*θ, τ_P_*) distribution contained two main density maxima, with extensions toward lower *θ* and toward higher *τ_P_* (Fig. 3b). Elliptical ROIs were defined within this distribution and the selected pixels were mapped back to the image (Fig. 3a).

Projecting the same pixel selections onto the native phasor plots revealed their distinct distributions in the two representations (Fig. 3c,d). The yellow and green populations over-lapped substantially in the spectral phasor plot but were more clearly separated in the lifetime phasor plot. Conversely, the purple and yellow populations were more clearly distinguished in the spectral representation than in the lifetime representation.

### Multimodal mapping across the Tau assembly series

We next analyzed Tau samples incubated for 1, 4, and 48 h. Pixels from the complete set of fields acquired at the three time points were pooled into a common (*θ, τ_P_*) distribution. The same elliptical ROIs were defined once on this pooled data set, without using the incubation time as a criterion, and subsequently propagated to the corresponding image pixels (Fig. 4). Although defined independently of incubation time, the different ROI populations were not equally represented across the series. The 1 h condensates were predominantly associated with the high-*θ*, shorter-*τ_P_* region of the distribution. At 4 h, the dominant populations shifted toward intermediate coordinates, whereas the 48 h assemblies showed a greater contribution from regions with longer *τ_P_* and lower *θ*. The distribution of ACDAN spectral–lifetime signatures thus shifted over the incubation series.

**Figure 4:**
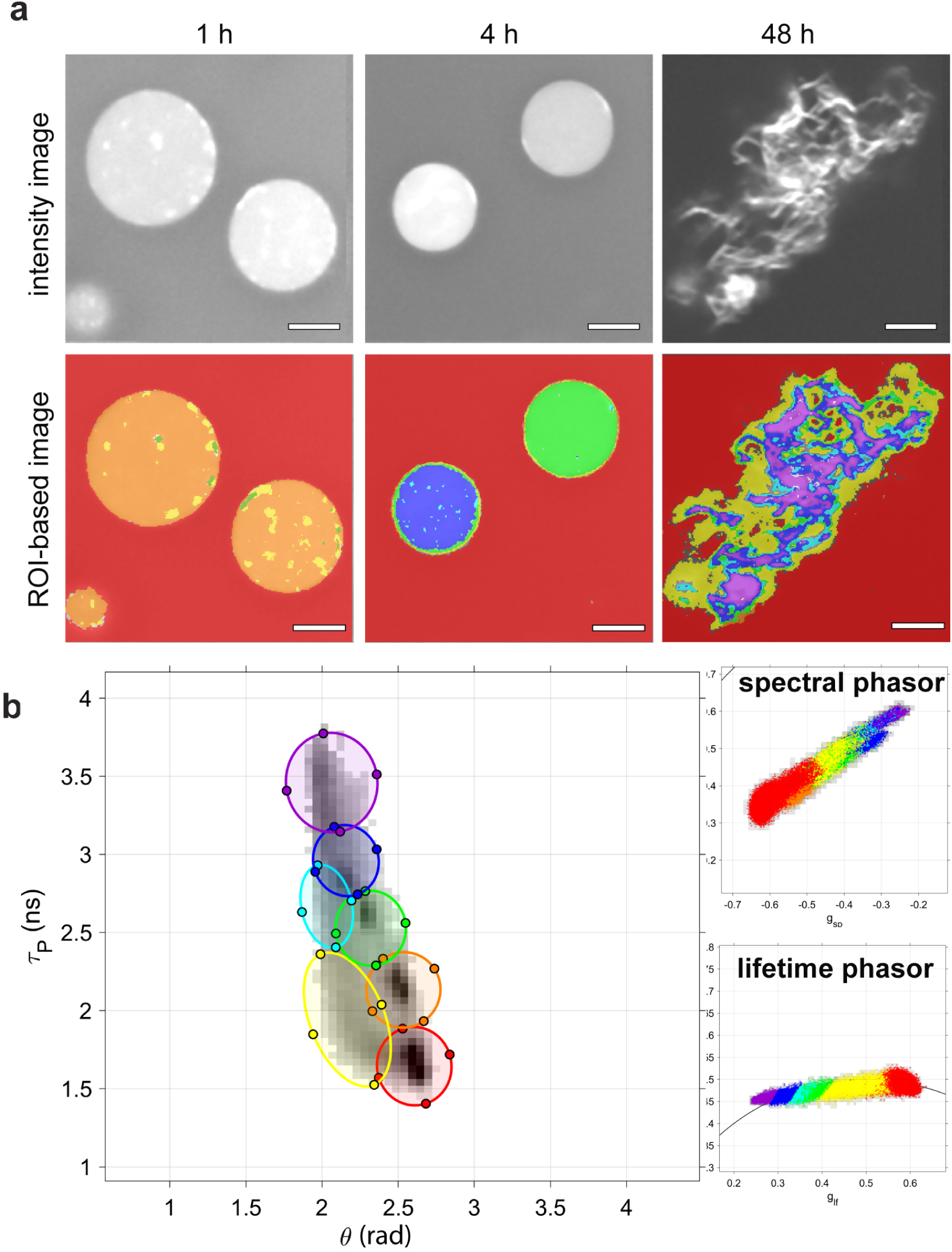
Multimodal mapping across the Tau assembly series. (a) Tau assemblies labeled with ACDAN at 1 h, 4 h and 48 h. Top row: hyperspectral intensity. Bottom row: pixel-wise assignment to the regions of interest defined in (b). (b) Intensity-weighted density in the combined (*θ, τ_P_*) plane, pooling all valid pixels from the three time points. Seven elliptical regions of interest defined on this distribution are propagated pixel-wise to each field in (a). Insets: spectral phasor (top) and lifetime phasor (bottom) of the same pixels, colored by ROI assignment, with the density of all valid pixels in grey. Scale bars: 5 µm.

The spatial organization of these populations also changed with time. At 1 h, the condensates were dominated by a common interior population while retaining localized spectral features similar to those described above. At 4 h, individual condensates remained comparatively homogeneous internally, but different condensates within the same field were assigned predominantly to different ROIs. Condensate-to-condensate heterogeneity was already detectable at this stage, even when each condensate had a relatively uniform internal signature. By 48 h, the round condensate morphology was replaced by an extended network containing several ROI populations distributed in spatially organized domains.

To assess whether the two coordinates provided interchangeable descriptions of the imaging data, each was used to predict the other by leave-one-field-out cross-validation. The linear models yielded 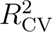 = 0.688 for *τ_P_* predicted from *θ* and 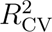 = 0.685 for *θ* predicted from *τ_P_*; allowing a cubic relationship did not improve either prediction (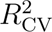 = 0.684 in both directions; Supplementary Note 2 and Supplementary Table S4). Thus, the two coordinates are strongly related, but neither fully reconstructs the other in this dataset.

### Thioflavin T and transmission electron microscopy of late-stage assemblies

To compare the ACDAN response with an established amyloid-sensitive probe, the same incubation series was analyzed independently using Thioflavin T (ThT, Supplementary Fig. S1b). Binding of ThT to amyloid-like cross-*β* structures strongly alters its fluorescence, particularly by restricting the intramolecular motions responsible for non-radiative relaxation.^37–39^ The ThT data were processed using the same multimodal workflow, with ROIs defined on the pooled (*θ, τ_P_*) distribution and propagated to the corresponding images (Fig. 5a,b).

**Figure 5:**
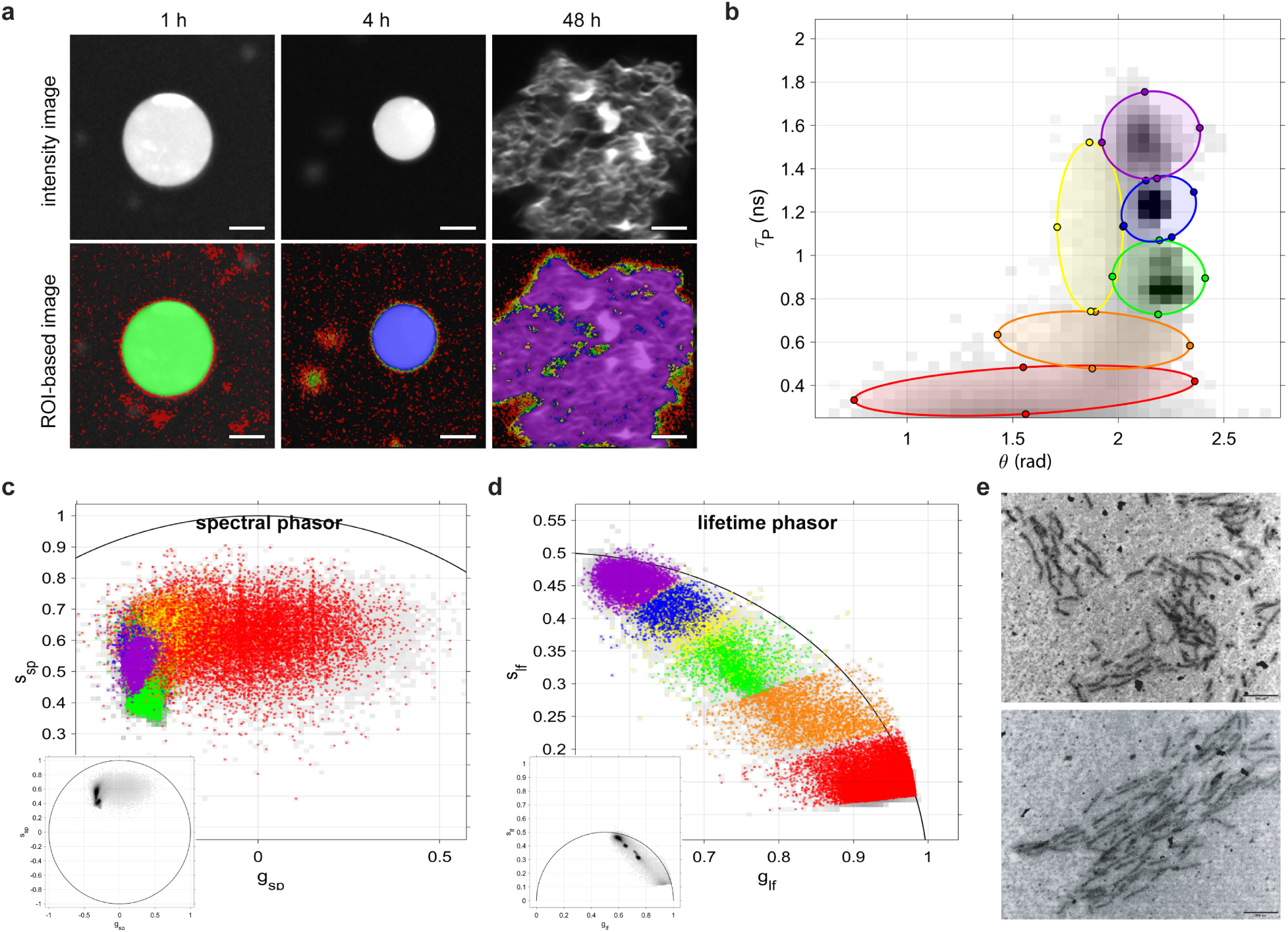
Thioflavin T and transmission electron microscopy of late-stage Tau assemblies. (a) Tau assemblies labeled with ThT at 1 h, 4 h and 48 h. Top row: hyper-spectral intensity. Bottom row: pixel-wise assignment to the regions of interest defined in (b). (b) Intensity-weighted density in the combined (*θ, τ_P_*) plane, pooling all valid pixels from the three time points, with six elliptical regions of interest defined on this distribution. (c) Spectral phasor and (d) lifetime phasor of the same pixels, colored by ROI assignment. Gray shading is the density of all valid pixels, and the black curves are the unit circle in (c) and the universal semicircle in (d). Insets show the full phasor space. (e) Representative transmission electron micrographs of Tau assemblies at 48 h. Scale bars: 5 µm (a) and 200 nm (e).

In contrast to ACDAN, the main separation between ThT populations occurred along the lifetime dimension. The distribution extended from short *τ_P_* values, prevalent at the earlier stages and in the surrounding phase, toward progressively longer lifetimes that became enriched in the 48 h structures. Changes in *θ* were comparatively smaller, and the populations showed substantial overlap along the spectral coordinate. The same behavior was apparent in the native phasor representations: ThT populations were ordered along the lifetime phasor trajectory (Fig. 5d), whereas they overlapped extensively in the spectral phasor plot (Fig. 5c). This difference was also reflected spatially. At 1 and 4 h, the condensates were each dominated by a single ROI and showed little internal differentiation with ThT. At 48 h, several lifetime-associated populations were distributed across the extended assemblies. Transmission electron microscopy performed at 48 h revealed elongated assemblies with protofibrillar-like morphology (Fig. 5e).

## Discussion

The spectral and lifetime responses of ACDAN differed in their sensitivity to the same changes in environment. This was evident in the reference solvents and became particularly useful in the Tau samples, where features that were clear in one coordinate were less apparent in the other. The shared (*θ, τ_P_*) plane is used here as a descriptive space alongside the native spectral and lifetime phasor plots, rather than as a new phasor domain, so that the same pixel populations can be followed across all representations.^12,18,19^ Leave-one-field-out prediction accounted for about 69% of the variance of either coordinate and was not improved by a cubic relationship (Supplementary Note 2), so *θ* and *τ_P_* behave as coupled but not interchangeable descriptors of this dataset. This comparison does not by itself imply that the two observables report distinct molecular variables.

ACDAN is a push–pull solvatochromic dye containing an electron-donating dimethylamino group and an electron-accepting carbonyl group (Supplementary Fig. S1a).^1^ Its spectral response reflects interactions with the surrounding medium and solvent relaxation following excitation, whereas its fluorescence lifetime depends on the combined rates of radiative and non-radiative deactivation. ^4^ Relative to glycerol and ethanol, the shorter lifetime and lower quantum yield in water were associated with a pronounced increase in the apparent non-radiative rate and a smaller decrease in the apparent radiative rate (Supplementary Table S2).

The cuvette measurements also show that the two observables do not order the solvents in the same way. Ethylene glycol and glycerol have emission maxima around 2 nm apart and the same apparent radiative rate, yet their apparent non-radiative rates differ by nearly a factor of two, while glycerol and ethanol have almost the same fluorescence lifetime and emission maxima separated by 20 nm (Supplementary Table S2). For PRODAN and related aminonaphthalenes, the position of the emission band has been shown to respond to the dipolarity and polarizability of the medium together with a specific hydrogen-bonding contribution, so that in protic solvents it does not report polarity alone. ^1,40,41^ Non-radiative deactivation has been linked to a different property of the medium, since in derivatives of this family designed to isolate it the quenching observed in protic solvents follows the hydrogen-bond donating ability of the solvent rather than its dipolarity or basicity.^42,43^ The present series is consistent with that division. The apparent non-radiative rate is an order of magnitude larger in water, the strongest hydrogen-bond donor of the four solvents, than in ethanol, and the two polyols fall between them while sharing almost the same emission maximum. The viscosity of the medium does not order the series, since glycerol and ethanol differ in it by orders of magnitude while their lifetimes differ by about 0.1 ns. The two observables also differ in how strongly they register these contributions. Because the apparent radiative rate lies between 0.24 and 0.25 ns*^−^*^1^ in the three organic solvents, the difference in apparent non-radiative rate between glycerol and ethanol shortens the fluorescence lifetime by only about 0.1 ns, whereas the same pair separates by 20 nm in emission maximum. In this solvent series, similar lifetimes need not correspond to similar spectral responses.

Because the ROIs in the incubation series were defined without reference to incubation time, the association between particular populations and stages emerged only after the selected pixels were mapped back to the corresponding samples. Differences between individual condensates at 4 h and coexisting populations within single assemblies at 48 h suggest that condensate-to-condensate differences precede the intra-assembly heterogeneity of the later stage.

Across the series, the spectral angle decreased and the phase lifetime increased between the early condensates and the 48 h assemblies (Fig. 4). Using the reference-solvent behavior as a qualitative guide, the decrease in *θ* together with the increase in *τ_P_*suggests a progressive restriction of dipolar relaxation in the environment sampled by ACDAN as the assemblies mature. Reduced dipolar relaxation inside condensates relative to the surrounding dilute phase has been described in several systems, including Tau condensates formed without aggregation-promoting cofactors.^25,27,28^ Independently, spatially-resolved NMR measurements of Tau condensates showed a reduction in water content in the dense phase relative to the dilute phase.^44^ These observations are compatible with an ACDAN-sensitive micro-environment that continues to evolve after condensate formation. At 48 h, the displacement in the ACDAN response coincided with the enrichment of longer-lived ThT populations and with the protofibrillar-like morphologies observed by TEM. A comparable increase in heterogeneity within individual condensates accompanied a reduction in molecular mobility during aging of *β*-lactoglobulin condensates,^35^ in which the spectral and lifetime readouts were obtained from separate probes rather than from a single probe in the same field of view. The origin of these changes cannot be deduced from ACDAN fluorescence alone. The maps describe spatial variations in fluorescence and do not establish whether the progressive shift reflects a continuous change throughout the dense phase or the emergence of structurally distinct regions during aging. Both behaviors have been described in aging condensates, including age-dependent concentration changes in PGL-3 condensates^45^ and liquid–solid coexistence during Tau condensate aging.^46^ The aqueous environment inside protein condensates differs from the surrounding dilute phase and can change with protein organization, ionic conditions, and condensate material properties, ^25,26,28^ and in Tau, changes in water structure have been associated with changes in condensate rheology under different ionic conditions. ^28^ Tau condensates also evolve differently depending on the conditions under which phase separation occurs.^30–32^ The changes detected here are therefore changes in the microenvironment accompanying assembly maturation under the conditions used in this study, not evidence for a universal condensate-to-fibril pathway. Heparin-induced Tau filaments are themselves polymorphic and differ from those found in Alzheimer’s and Pick’s diseases,^47^ so the late-stage assemblies examined here are not assumed to be structurally representative of disease filaments, although their protofibrillar-like morphology is consistent with progression toward aggregated structures at the latest time point.

ACDAN and ThT respond to different aspects of their local environment. ACDAN is sensitive to solvation and solvent dipolar relaxation, whereas ThT fluorescence is strongly affected by its binding environment and the restriction of intramolecular motion. ^3,37,39^ The spatial differences detected with ACDAN in early condensates do not establish earlier detection of amyloid formation. For ThT, changes in fluorescence lifetime can reflect differences in binding environment and local rigidity, while self-quenching at high local dye concentrations can also shorten the lifetime.^38,39^ The longer-lived populations observed at 48 h are thus consistent with changes in the environments experienced by ThT, but do not identify a particular Tau conformation.

Linear combination rules apply to Cartesian phasor coordinates under defined conditions and do not transfer to derived variables such as *θ* and *τ_P_*,^12,18,19,48^ so the combined plane is not used for unmixing. The elliptical ROIs select populations with different fluorescence signatures without assigning them to molecular states, and they serve as a fixed reference that is propagated pixel-wise to the spectral phasor, the lifetime phasor, and the image without being redefined in each domain.

The same workflow can be applied to other environment-sensitive probes used in condensates, including Nile Red and viscosity-sensitive BODIPY molecular rotors, ^49^ and to cellular samples, in which ACDAN has already been used. ^10^

### Conclusions

The spectral and lifetime responses of ACDAN distinguished the reference solvents in different ways, and in the imaging data neither coordinate fully predicted the other. Relating the spectral phase angle *θ* and the phase lifetime *τ_P_* in a shared plane, while keeping the native phasor representations and the pixel-wise link to the image, allowed both responses to be examined together. In Tau, this revealed differences among individual condensates at 4 h and heterogeneous domains within the extended 48 h assemblies, together with a decrease in *θ* and an increase in *τ_P_* across the incubation series. Thioflavin T analyzed in the same way separated mainly along the lifetime coordinate, showing that the information recovered from the combined plane depends on the photophysics of the probe.

## Methods

### Phasor analysis: general framework

The phasor approach represents fluorescence signals in a two-dimensional plane using the sine and cosine components of their Fourier transform at a selected harmonic frequency. The transformation itself does not require fitting the fluorescence signal to a predefined model. In their native Cartesian representation, phasor coordinates also preserve the linear combination of intensity-weighted fluorescent components. The same mathematical formalism can be applied to emission spectra and fluorescence decays, although the input signal and the physical interpretation of the resulting coordinates differ between the two domains.^12,18,19^

#### Spectral phasor

Emission spectra were recorded in *N* contiguous detection channels of equal width Δ*λ*, with center wavelengths *λ*_1_*,…, λ_N_*. For each pixel spectrum *I*(*λ_k_*), the first-harmonic spectral phasor coordinates were calculated as^19^

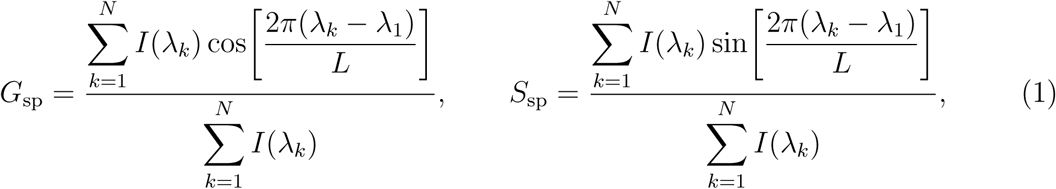

where *L* = *N* Δ*λ* is the period of the transform. For the detection window used here, *L* = 31 × 8.9 = 275.9 nm.

The spectral phase angle was obtained using the four-quadrant inverse tangent,

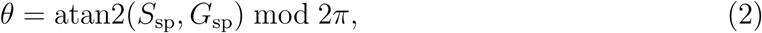

and the spectral modulus was calculated as 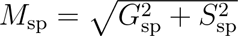. Within a fixed acquisition window, changes in *θ* follow displacement of the spectral distribution, whereas *M*_sp_ contains information related to spectral width. ^19^ For small displacements, an angular difference can be expressed as an approximate wavelength-equivalent displacement,

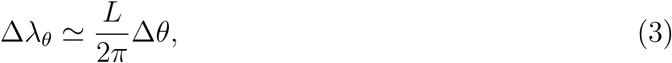

which corresponds to 43.9 nm per radian for the acquisition window used here. This conversion is used only to provide an intuitive scale for differences in *θ* and should not be interpreted as an independent determination of the spectral center of mass.

No correction for the wavelength-dependent sensitivity of the spectral detector was applied before the phasor transformation. All comparisons were therefore performed using the same detector configuration and spectral window. Consequently, absolute *θ* values are specific to this acquisition configuration, whereas comparisons among samples measured under the same conditions remain internally consistent.

#### Lifetime phasor

In the lifetime domain, raw phasor coordinates were computed from the fluorescence intensity decay histogram *I*(*t_j_*) recorded at laser repetition frequency *f*, with angular frequency *ω* = 2*πf*, as^11,18^

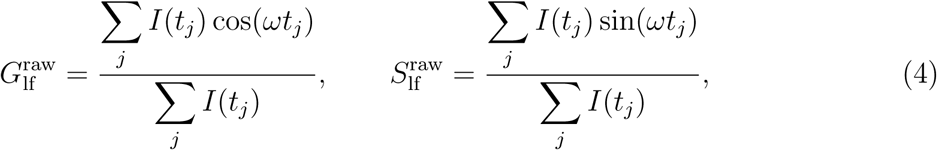

where the sums run over the time channels *j* spanning one full pulse period. These raw coordinates were then transformed into calibrated coordinates (*G*_lf_ *, S*_lf_) using an external reference of known lifetime, as described below.

Two lifetime descriptors follow directly from the calibrated coordinates without a fitting model. The phase lifetime is

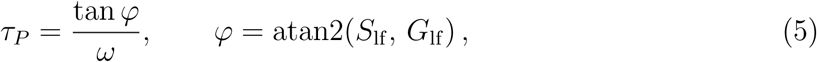

and the modulation lifetime is

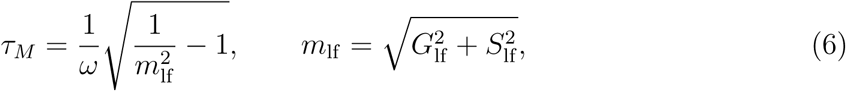

reflecting respectively the phase shift and the demodulation of the emission relative to the excitation.^11,12,18,21^ The modulation is written *m*_lf_ to distinguish it from the spectral modulus *M*_sp_ of Eq. 1.

For an ideal monoexponential decay of lifetime *τ*, both *τ_P_* and *τ_M_* equal *τ*, and the phasor lies on the universal semicircle of radius 1*/*2 centered at (1*/*2, 0). For a signal composed of a non-interacting mixture of positive exponential decay components, the phasor lies within the semicircle and, for an interior point, *τ_P_ < τ_M_*.^4,12,18^ The ratio

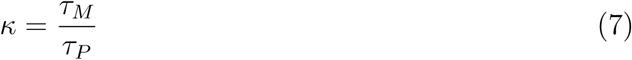

equals unity for an ideal monoexponential decay and exceeds unity for an interior phasor under this mixture model. Although *κ* is calculated without fitting a decay model, it is not a unique measure of heterogeneity: its value also depends on the component lifetimes, their relative intensity contributions, and the modulation frequency. ^11,12^ Phasors outside the universal semicircle are not interpreted using this positive-mixture model. Their handling is described under *Lifetime phasor calibration*.

### Protein expression and purification

Full-length 2N4R human Tau was expressed recombinantly in *Escherichia coli* BL21(DE3) CodonPlus RIL cells (Agilent, Santa Clara, USA) transformed with the plasmid tau/pET29b (Addgene #16316). Briefly, 20 ng of plasmid was added to 20 µL of chemically competent cells and subjected to heat shock at 42 °C for 30 s. After recovery in LB medium at 37 °C for 60 min, transformed bacteria were plated on LB agar supplemented with 50 µg/mL kanamycin and incubated overnight at 37 °C. Single colonies were grown overnight at 37 °C and 200 rpm in 25 mL LB broth containing 50 µg/mL kanamycin, then inoculated 1:100 into Terrific Broth (Carl Roth, Karlsruhe, Germany) with 50 µg/mL kanamycin and incubated at 37 °C until OD_600_ = 0.6. Protein expression was induced with 1 mM isopropyl *β*-D-1-thiogalactopyranoside (IPTG; Carl Roth) for 3 h at 37 °C.

For purification, the bacterial pellet was resuspended in buffer A (20 mM PIPES pH 6.3, 15 mM NaCl, 1 mM EDTA, 1 mM MgSO_4_, 1 mM DTT) and lysed by ultrasonication (SFX250 Digital Sonifier, Branson) on ice for 10 min (0.5 s pulses, 70 W, 60% amplitude). After centrifugation at 10,000 g and 4 °C, NaCl was adjusted to 500 mM and the supernatant was boiled for 10 min to precipitate folded bacterial proteins. After a second centrifugation, the supernatant was diluted to 80 mM NaCl with buffer A and applied to a cation exchange column (HiTrap Capto SP ImpRes 5 mL; Cytiva) on an ÄKTA Pure 25 FPLC system (Cytiva). Tau was eluted with a linear NaCl gradient. Fractions containing Tau (eluting at approximately 1.5 M NaCl) were pooled, concentrated using a 10 kDa centrifugal filter (Sartorius AG, Göttingen, Germany), and stored in aliquots at −80 °C. Purity was confirmed by SDS-PAGE. The purification protocol was adapted from Ferrari and Rüdiger. ^50^

### Preparation of Tau condensates and assemblies

Purified 2N4R Tau was diluted to 25 µM in assembly buffer (10 mM HEPES pH 7.4, 10 mM NaCl, 1 mM EDTA, 15% w/v polyethylene glycol 8000 (PEG-8000)) and heparin was added at a 4:1 Tau:heparin molar ratio (6.25 µM heparin). PEG-8000 served as a macromolecular crowding agent to promote condensate formation, and heparin acted as a polyanion co-factor for Tau assembly, following established protocols. ^32,50^ Samples were incubated at 37 °C without agitation and aliquots were collected from the same preparation at 1 h, 4 h, and 48 h to follow assembly maturation.

### Fluorescent probes and sample preparation for microscopy

ACDAN and Thioflavin T (ThT) were dissolved in dimethyl sulfoxide (DMSO) to prepare stock solutions of 5 mM (ACDAN) and 3 mM (ThT). Working solutions were prepared by dilution into the Tau assembly reaction immediately before imaging, yielding a final probe concentration of 5 µM for both probes and a final DMSO content of 0.1% (v/v) for ACDAN and 0.17% (v/v) for ThT. ACDAN and ThT were used in separate experiments. Probes were allowed to equilibrate with the sample for approximately 5 min before image acquisition. For microscopy, 20 µL aliquots were deposited into µ-Slide 15 Well 3D chambers (ibidi GmbH, Gräfelfing, Germany). Chamber surfaces were passivated prior to use by incubation with 1 mg/mL poly-L-lysine-*graft*-polyethylene glycol (PLL-*g*-PEG; SuSoS AG, Dübendorf, Switzerland) to minimize non-specific protein adsorption.

### Photophysical characterization of ACDAN in reference solvents

Absolute quantum yields (Φ) and fluorescence lifetimes (*τ*_exp_) of ACDAN were determined in water, ethylene glycol, glycerol, and ethanol. Full instrumental details, sample preparation and the analysis of the absorption and emission spectra are given in the Supplementary Methods and Supplementary Fig. S2. For a monoexponential fluorescence decay, the lifetime is related to the radiative and non-radiative rates by

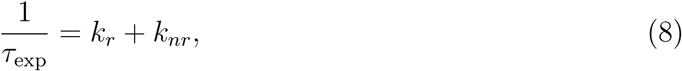

where *k_r_* and *k_nr_* are the apparent radiative and non-radiative rates derived under the monoexponential approximation. Using the absolute quantum yield, these quantities were calculated as

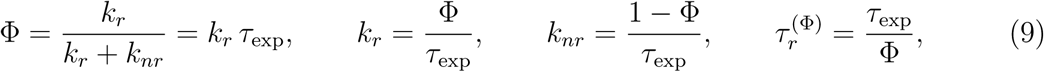

following standard photophysical definitions.^4^ The Strickler–Berg calculation and the resulting radiative-lifetime estimates are presented in Supplementary Note 1 and Supplementary Table S3.

### Hyperspectral and FLIM microscopy

Multimodal imaging was performed at the Advanced Optical Microscopy Centre (AOMC) of Hasselt University using a Zeiss LSM 880 NLO confocal microscope. Samples were excited at 810 nm via two-photon absorption using a pulsed Ti:Sapphire Mai-Tai-HPDS laser and a water-immersion C-ApoChromat 63×/1.20 W Korr objective. The average excitation power at the sample was not quantified in absolute terms, but was kept constant across all acquisitions reported here, together with detector gain and integration time, so that intensities can be compared between samples. Imaging was performed at room temperature, approximately 22 °C.

#### Field-of-view matching

FLIM and hyperspectral data were acquired sequentially from the same field of view without moving the microscope stage or objective between acquisitions. FLIM was acquired first and required approximately 5 min per field, followed immediately by the hyperspectral lambda scan, which required approximately 10 s. Both acquisitions used the same pixel grid, allowing direct pixel-wise correspondence between the two datasets. Control measurements acquired in the reverse order produced comparable spectral and life-time distributions.

#### Hyperspectral imaging

Hyperspectral imaging was performed in lambda-scan mode using the 32-channel GaAsP spectral detector (Quasar unit; Zeiss), with the confocal pinhole set to 1 Airy unit. Emission was collected in *N* = 31 channels with center wavelengths from *λ*_1_ = 423.1 to *λ*_31_ = 690.1 nm, corresponding to a spectral sampling of Δ*λ* = 8.9 nm per channel. These values define the period *L* of the spectral phasor transform in Eq. 1. Images were acquired at 256 × 256 pixels with a pixel size of 110 nm, corresponding to a field of view of 28.2 µm, and a pixel dwell time of approximately 1.5 µs.

#### Fluorescence lifetime imaging

FLIM was acquired in the BiG.2 non-descanned detection unit using a single GaAsP detection channel equipped with a 550/200 nm bandpass emission filter, so no combination of channels was required before the phasor transform. Photons were time-tagged by a PicoQuant MultiHarp 150 multichannel TCSPC device and acquisition was controlled via SymphoTime64 software (PicoQuant GmbH, Berlin, Germany) at a laser repetition rate of 80 MHz, corresponding to *ω* = 2*π* ×80 ×10^6^ rad s*^−^*^1^ and a 12.5 ns pulse period. Decays were histogrammed into 250 time channels of 50 ps. Acquisition was continued until at least 500 photons had accumulated per pixel in homogeneous fields, or per pixel within the structures of interest in heterogeneous samples.

Note that the two modalities do not sample identical spectral windows: *θ* is computed over the full 423–690 nm detection range, whereas *τ_P_* is measured over the 450–650 nm band defined by the FLIM emission filter. Both windows encompass the main emission band of ACDAN in all solvents examined.

Both datasets were pre-processed using PAM-PIE,^51^ which computed the spectral and FLIM phasor coordinates of each pixel according to Eqs. 1–6. The two modalities were processed independently. Their coordinate and intensity maps were then combined by corresponding image coordinates into a single structure per field of view. These combined structures are the input to the multimodal analysis described below. Experiments were performed on independent sample preparations, and representative fields are shown in all figures.

#### Display conventions

For the reference solvents, the wavelength and *τ_P_* maps use limits shared across all samples so that colors are directly comparable between panels. Intensity images are displayed with per-panel limits because the dynamic range differs by almost an order of magnitude across the solvent series, and the corresponding mean pixel intensities are stated in the text. The *τ_P_*color ramp is deliberately reversed with respect to the usual FLIM convention, placing warm colors at short lifetimes, so that a sample in which a short lifetime coincides with a red-shifted emission retains a consistent color in both of its maps. All color scale limits are shown in the corresponding figures.

### Lifetime phasor calibration

Lifetime phasor coordinates were calibrated against an external reference of known monoexponential decay, consisting of 100 nM ATTO 488 in a saturated aqueous potassium iodide solution diluted 1:1000 in Milli-Q water. The reference was measured on the same microscope, under the same acquisition settings as the samples, at the same repetition rate, with the same number of time channels and using the same detection channel and emission filter. Its lifetime was determined in situ by tail fitting of the spatially integrated decay, giving *τ*_ref_ = 3.59 ns. Because tail fitting is independent of the phase and modulation calibration being derived from it, this procedure is not circular, and determining *τ*_ref_ on the imaging system itself removes any systematic difference between the reference and sample measurements.

The theoretical phasor coordinates of the reference are

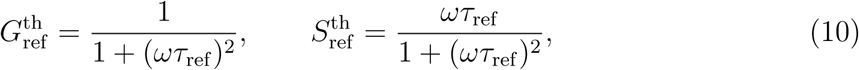

from which a phase offset Δ*φ* = *φ*^th^ − *φ*^raw^ and a modulation factor *c* = *m*^th^*/m*^raw^ were obtained, where *φ*^raw^ and *m*^raw^ are the phase and modulus of the measured reference phasor. Sample coordinates were then calibrated by rotation and scaling,

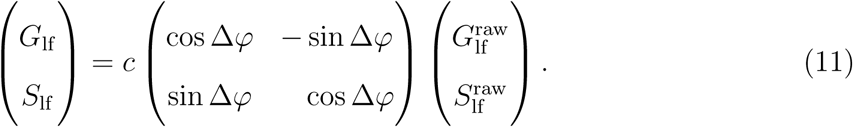

A single pair of calibration parameters was determined per acquisition session and applied to all pixels of all samples acquired in that session. No sample-derived rotation was applied at any point, so the position of each sample relative to the universal semicircle is determined solely by the external reference.

The instrument response function was measured separately using 100 nM ATTO 488 in saturated potassium iodide, in which the fluorescence lifetime is quenched to a value short compared with the instrument timing resolution. The IRF was used to verify the timing stability of the imaging system, but not for the phasor transform, which requires only an external lifetime reference.

Pixels whose calibrated coordinates fell marginally outside the universal semicircle were retained rather than clipped. For these pixels, *τ_P_* is reported only as a phase-derived descriptor under the acquisition and calibration conditions used. It is not interpreted as a unique molecular lifetime. An outside-circle position can reflect excited-state processes, including time-dependent solvent relaxation, or measurement and processing effects. The phasor position alone does not establish which explanation applies. ^12^ For these pixels, *τ_M_* and *κ* are not interpreted using the standard positive-mixture model.

### Pixel-wise derived maps and decay analysis

In addition to the phasor coordinates, several pixel-wise and field-integrated quantities are reported for the reference measurements. These were computed on pixels considered valid, meaning those exceeding 20 counts in the summed hyperspectral image for spectral quantities, and 100 photons in the FLIM acquisition with a phase lifetime within the physically plausible range 0.25 to 5.5 ns for lifetime quantities.

#### Wavelength of maximum emission

For each pixel, the emission maximum *λ*_max_ was estimated from the highest-intensity spectral channel, with log-parabolic interpolation using its two neighboring channels for interior peaks.

#### Mean photon arrival time

For each pixel, the mean photon arrival time was computed directly from the raw decay histogram as

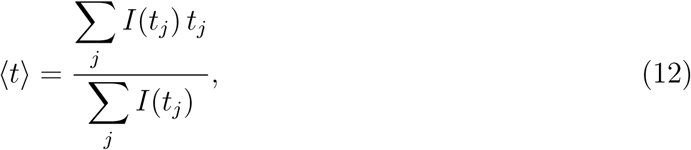

without deconvolution or offset correction. This quantity therefore contains the instrument response offset and is not a fluorescence lifetime.

#### Pixel-averaged emission spectra

Spectra shown for imaging data are pixel-averaged across all valid pixels in a field. When spectra from different samples are overlaid, each curve is divided by its own integral over the detection window, so that each spectrum encloses a unit area and the comparison is of band position and shape rather than amplitude. This is also the normalization applied internally by the spectral phasor transform, in which *G* and *S* are ratios to the integrated intensity. Peak positions and full widths at half maximum were obtained by fitting a Gaussian function to the averaged spectrum before normalization, and the uncertainty quoted for the peak position is the standard error of that fit.

#### Spatially integrated decay analysis of imaging data

Spatially integrated decays were obtained by summing the TCSPC histograms of all valid pixels of a field. Decays were analyzed by tail fitting with a sum of *n* exponential components over a window starting 2.5 ns after the pulse, chosen to exclude the rising edge, where the signal is dominated by the instrument response. Poisson weighting was used, and weighted residuals are reported as 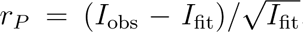. The number of components was selected using the Bayesian information criterion,

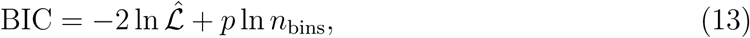

where L̂ is the maximized Poisson likelihood, *p* the number of free parameters and *n*_bins_ the number of fitted time channels. The constant offset returned by this fit was subtracted from the FLIM histograms before the phasor transform of Eq. 4. Individual pixel decays were not fitted for the phasor calculation. The fit was used only to estimate a field-level background offset and to characterize the spatially integrated decay.

### Multimodal feature-space construction and analysis

Before any masking or pooling, a 3 × 3 median filter was applied to the coordinate and intensity maps, as is customary in phasor analysis to reduce pixel-level noise without displacing cluster positions. ^12,18^ A pixel was considered valid when the spectral phasor angle, the phase lifetime, and the integrated intensities of both acquisitions were all defined, and when both the phase and modulation lifetimes fell within the physically plausible range 0.25 to 5.5 ns. Unless stated otherwise, every pooled distribution, region of interest, and derived quantity reported for Tau assemblies was computed on valid pixels so defined.

The spectral phasor angle *θ* and the phase lifetime *τ_P_* were combined into a joint (*θ, τ_P_*) state space, where each pixel contributes a single point whose coordinates encode both its spectral and temporal fluorescence characteristics. To construct the two-dimensional intensity-weighted histogram in this space, each pixel was assigned a composite weight based on the relative intensities of both modalities,

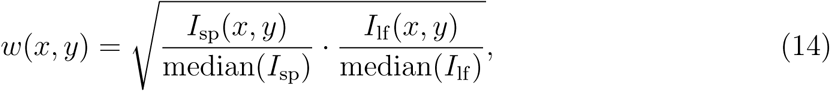

where *I*_sp_ and *I*_lf_ are the integrated intensities of the hyperspectral and FLIM acquisitions, respectively, and the medians are computed over all valid pixels in the field of view. The weight is the geometric mean of the two median-normalized intensities. It scales with their product, so a high intensity in one modality can compensate for a low intensity in the other. We therefore treat *w*(*x, y*) as a weighting choice for constructing the joint density, not as evidence that each included pixel has high photon counts in both modalities. The histogram was accumulated on a 120 × 120 grid and normalized to [0, 1] for display.

For heterogeneous samples, a global reference distribution was constructed by pooling valid pixels from all samples into a single (*θ, τ_P_*) histogram. Regions of interest were defined as ellipses drawn interactively on this pooled distribution, so that the same reference frame was shared across all datasets. Once defined, ROI boundaries were not redefined in any subsequent domain: the pixel sets selected in the combined (*θ, τ_P_*) plane were propagated without re-thresholding to the spectral phasor domain, to the lifetime phasor domain, and back to image space, maintaining a one-to-one linkage by image coordinate across representations. The analysis of Tau assemblies was implemented in MATLAB (The MathWorks, Natick, USA), and the reference-solvent and single-sample panels in Python.

### Cross-validated comparison of spectral and lifetime coordinates

To assess whether *θ* and *τ_P_* provided interchangeable descriptions of the imaging data, we used bidirectional leave-one-field-out cross-validation on the per-pixel coordinate data, independently of the region-of-interest selection used for display. In each fold, all pixels from one field of view were withheld, a least-squares linear relationship between the two coordinates was fitted using the remaining fields, and the withheld coordinate was predicted from the other. The procedure was repeated with predictor and response exchanged. Keeping each field intact during cross-validation prevents pixels from the same image from contributing to both model fitting and evaluation, following the general principle of separating dependent groups during predictive validation.^52,53^

Prediction performance was summarized from the concatenated out-of-fold predictions using the cross-validated coefficient of determination, *R*^2^, and the root-mean-square error (RMSE). The complete procedure was also repeated with a third-order polynomial to test whether a simple smooth nonlinearity improved reconstruction. Further details and the resulting values are given in Supplementary Note 2 and Supplementary Table S4.

### Transmission electron microscopy

Transmission electron microscopy was used to examine the morphology of Tau assemblies after 48 h. A 5 µL aliquot of the probe-free sample was applied to a formvar-coated carbon grid (200 mesh; Ted Pella Inc.) and allowed to adsorb for 60 s. Excess liquid was removed by blotting, after which 3 µL of 2% uranyl acetate was applied for 30 s. The grid was blotted again and air-dried.

Samples were imaged using a JEOL JEM-1400 Flash transmission electron microscope operated at 80 kV. Images were recorded with an Emsis Xarosa CMOS camera using RADIUS 2.1 software. Fields were examined over a range of magnifications, and representative images are shown. Protofibrillar-like morphologies were observed in independent sample preparations.

### Supporting Information Available

Additional photophysical characterization methods; Strickler–Berg analysis; cross-validated comparison of spectral and lifetime coordinates; chemical structures of ACDAN and Thioflavin T; absorption spectra in the reference solvents; quantum-yield, photophysical, Strickler–Berg, and cross-validation tables (PDF).

## Acknowledgement

During preparation of this work, the authors used Claude (Anthropic) to assist with implementing and debugging parts of the analysis and figure-generation code, and ChatGPT (OpenAI) for language editing. The authors critically reviewed all AI-assisted output and take full responsibility for the content of the publication.

We acknowledge the Advanced Optical Microscopy Centre at Hasselt University for support with the microscopy experiments. Microscopy was made possible by the Research Foundation Flanders (FWO, project G0H3716N). Transmission electron microscopy was made possible by the Research Foundation Flanders (FWO, grant I000220N). We thank Dr. Sam Duwe for his skillful technical assistance with the optical microscopy experiments, and Marc Jans and Wendy Vandendries for their skillful technical assistance with transmission electron microscopy. We also thank the members of the Dynamic Bioimaging Lab for stimulating scientific discussions throughout the development of this work. We thank Prof. Dr. Wouter Maes (UHasselt, IUMAT, DSOS) for providing access to the absorption and fluorescence spectroscopy instrumentation used for the photophysical characterization and for his critical reading of the manuscript.

This work was supported by the Research Foundation Flanders (FWO) through the INSITE project (FWO, G0A8L24N) to J.H. and E.C. S.B. acknowledges financial support from UHasselt and the FWO (G0D1521N).

## Author contributions

L.C.R. designed the study, performed the microscopy and spectroscopy experiments, developed the analysis framework, and wrote the manuscript. Y.V. contributed to sample preparation, protein production, and microscopy experiments. S.B. contributed to cuvette photophysical characterization. I.D. and E.C. contributed to the writing and revision of the manuscript, and E.C. acquired funding. J.H. supervised the study, acquired funding, and revised the manuscript. All authors read and approved the final version.

## Data availability

The datasets generated and analyzed during this study are not publicly available because of the size of the raw time-resolved and hyperspectral acquisitions, but are available from the corresponding author upon reasonable request.

## Code availability

The MATLAB implementation used to generate the multimodal phasor representations of Tau assemblies and the Python implementation used for the reference-solvent and single-sample panels are available at https://gitlab.com/leandro.cruzrodrigez/multid-phasors, released under the MIT license.

---

**TOC Graphic.**
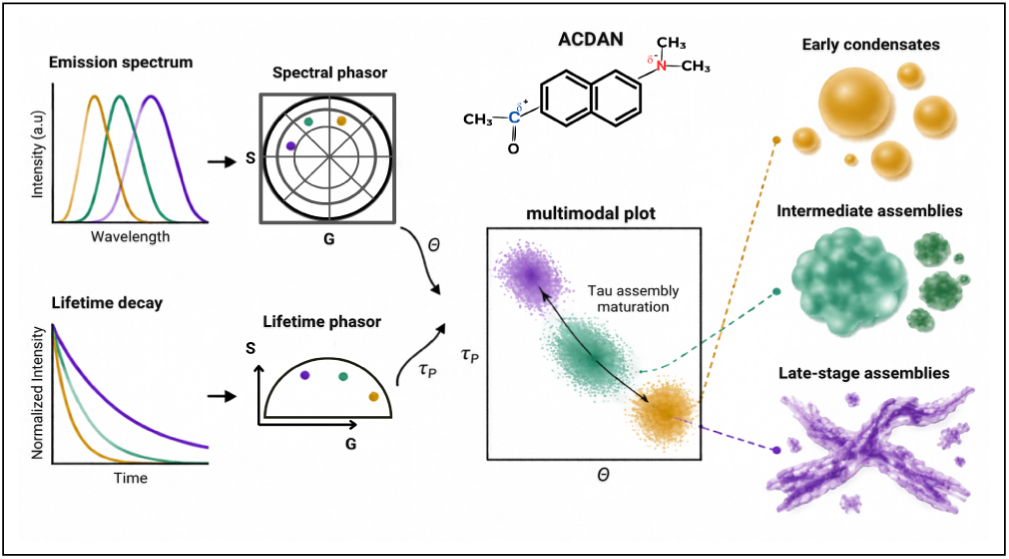

## References

(1) Weber, G.; Farris, F. J. Synthesis and spectral properties of a hydrophobic fluorescent probe: 6-propionyl-2-(dimethylamino) naphthalene. Biochemistry 1979, 18, 3075–3078.

(2) Parasassi, T.; Krasnowska, E. K.; Bagatolli, L.; Gratton, E. Laurdan and Prodan as polarity-sensitive fluorescent membrane probes. Journal of fluorescence 1998, 8, 365– 373.

(3) Otaiza-González, S.; Cabadas, M.; Robert, G.; Stock, R. P.; Malacrida, L.; Lascano, R.; Bagatolli, L. A. The innards of the cell: studies of water dipolar relaxation using the ACDAN fluorescent probe. Methods and Applications in Fluorescence 2022, 10, 044010.

(4) Lakowicz, J. R. Principles of Fluorescence Spectroscopy, 3rd ed.; Springer: New York, 2006.

(5) Bagatolli, L. A. Monitoring membrane hydration with 2-(dimethylamino)-6-acylnaphtalenes fluorescent probes. Membrane Hydration: The Role of Water in the Structure and Function of Biological Membranes 2015, 105–125.

(6) Mitra, A. K.; Das, S. A Journey Through Advances in 6-Acyl-2-(Dimethylamino) Naphthalene (DAN) Fluorescent Probes: Synthesis, Photophysical Insights, and Beyond. Journal of Molecular Structure 2025, 142817.

(7) Cho, Y. H. A.; Sandoval-Altamirano, C.; Sánchez, S. A.; Pizarro, N.; Romero-Parra, J.; Coeffard, V.; Gunther, G. An overlooked view at Weber’s DAN solvatochromic probes: photosensitization and photostability. Journal of Photochemistry and Photobiology A: Chemistry 2026, 474, 116994.

(8) De Luca, G.; Ferrara, V.; Pignataro, B.; Vetri, V.; Sancataldo, G. A Fluorescent Perspective on Water Structuring: ACDAN in Salt Solutions and Hydrogels. Biophysica 2024, 4, 619–633.

(9) Vorontsova, I.; Vallmitjana, A.; Torrado, B.; Schilling, T. F.; Hall, J. E.; Gratton, E.; Malacrida, L. In vivo macromolecular crowding is differentially modulated by aquaporin 0 in zebrafish lens: Insights from a nanoenvironment sensor and spectral imaging. Science Advances 2022, 8, eabj4833.

(10) Thoke, H. S.; Thorsteinsson, S.; Stock, R. P.; Bagatolli, L. A.; Olsen, L. F. The dynamics of intracellular water constrains glycolytic oscillations in Saccharomyces cerevisiae. Scientific Reports 2017, 7, 16250.

(11) Ranjit, S.; Malacrida, L.; Jameson, D. M.; Gratton, E. Fit-free analysis of fluorescence lifetime imaging data using the phasor approach. Nature protocols 2018, 13, 1979–2004.

(12) Malacrida, L.; Ranjit, S.; Jameson, D. M.; Gratton, E. The phasor plot: a universal circle to advance fluorescence lifetime analysis and interpretation. Annual review of biophysics 2021, 50, 575–593.

(13) Recoulat Angelini, A. A.; Malacrida, L.; González Flecha, F. L. Fluorescence phasor analysis: basic principles and biophysical applications. Biophysical Reviews 2025, 17, 395–408.

(14) Weber, G. Resolution of the fluorescence lifetimes in a heterogeneous system by phase and modulation measurements. The Journal of Physical Chemistry 1981, 85, 949–953.

(15) Jameson, D. M.; Gratton, E.; Hall, R. D. The measurement and analysis of heterogeneous emissions by multifrequency phase and modulation fluorometry. Applied spectroscopy reviews 1984, 20, 55–106.

(16) Clayton, A.; Hanley, Q.; Verveer, P. Graphical representation and multicomponent analysis of single-frequency fluorescence lifetime imaging microscopy data. Journal of microscopy 2004, 213, 1–5.

(17) Redford, G. I.; Clegg, R. M. Polar plot representation for frequency-domain analysis of fluorescence lifetimes. Journal of fluorescence 2005, 15, 805–815.

(18) Digman, M. A.; Caiolfa, V. R.; Zamai, M.; Gratton, E. The phasor approach to fluorescence lifetime imaging analysis. Biophysical journal 2008, 94, L14–L16.

(19) Fereidouni, F.; Bader, A. N.; Gerritsen, H. C. Spectral phasor analysis allows rapid and reliable unmixing of fluorescence microscopy spectral images. Optics express 2012, 20, 12729–12741.

(20) Coucke, Q.; Parveen, N.; Fernández, G. S.; Qian, C.; Hofkens, J.; Debyser, Z.; Hendrix, J. Particle-based phasor-FLIM-FRET resolves protein-protein interactions inside single viral particles. Biophysical Reports 2023, 3, 100122.

(21) Malacrida, L.; Jameson, D. M.; Gratton, E. A multidimensional phasor approach reveals LAURDAN photophysics in NIH-3T3 cell membranes. Scientific reports 2017, 7, 9215.

(22) Scipioni, L.; Rossetta, A.; Tedeschi, G.; Gratton, E. Phasor S-FLIM: a new paradigm for fast and robust spectral fluorescence lifetime imaging. Nature methods 2021, 18, 542–550.

(23) Haas, K. T.; Fries, M. W.; Venkitaraman, A. R.; Esposito, A. Single-cell biochemical multiplexing by multidimensional phasor demixing and spectral fluorescence lifetime imaging microscopy. Frontiers in Physics 2021, 9, 637123.

(24) Shin, Y.; Brangwynne, C. P. Liquid phase condensation in cell physiology and disease. Science 2017, 357, eaaf4382.

(25) Mangiarotti, A.; Siri, M.; Tam, N. W.; Zhao, Z.; Malacrida, L.; Dimova, R. Biomolecular condensates modulate membrane lipid packing and hydration. Nature Communications 2023, 14, 6081.

(26) De Luca, G.; Malacrida, L.; Vetri, V.; Sancataldo, G. Unveiling water ordering in liquid–liquid phase separation using bovine serum albumin-polyethylene glycol systems. Journal of Molecular Liquids 2025, 433, 127865.

(27) Cruz Rodríguez, L.; Foressi, N. N.; Celej, M. S. Tracking protein transitions through fluorescence spectral phasor analysis with ACDAN. Biophysical Reports 2025, 5, 100209.

(28) Foressi, N. N.; Cruz Rodríguez, L.; Wilke, N.; Celej, M. S. Cation-Driven Modulation of Tau Condensates: Insights into Liquid–Liquid Phase Separation and Rheological Properties. Biomacromolecules 2025, 26, 3605–3616.

(29) Sabri, E.; Mangiarotti, A.; Dimova, R. Fluorescence-based mapping of condensate dielectric permittivity uncovers hydrophobicity-driven membrane interactions. Nature Communications 2026, 17, 3155.

(30) Kanaan, N. M.; Hamel, C.; Grabinski, T.; Combs, B. Liquid-liquid phase separation induces pathogenic tau conformations in vitro. Nature Communications 2020, 11, 2809.

(31) Chakraborty, P.; Zweckstetter, M. Phase separation of the microtubule-associated protein tau. Essays in Biochemistry 2022, 66, 1013–1021.

(32) Boyko, S.; Surewicz, K.; Surewicz, W. K. Regulatory mechanisms of tau protein fibrillation under the conditions of liquid–liquid phase separation. Proceedings of the National Academy of Sciences 2020, 117, 31882–31890.

(33) Mahendran, T. S.; Singh, A.; Srinivasan, S.; Jennings, C. M.; Neureuter, C.; Gindra, B. H.; Parekh, S. H.; Banerjee, P. R. Decoupling phase separation and fibrillization preserves activity of biomolecular condensates. Nature Communications 2026, 17, 2841.

(34) Mangiarotti, A.; Ambroggio, E. E.; Celej, M. S.; Malacrida, L. Unveiling biomolecular condensate physiology and pathology using spectral and time-resolved fluorescence microscopy of DAN probes. Biophysics Reviews 2026, 7, 041301.

(35) Anselmo, S.; Scollo, F.; Sancataldo, G.; Lanzanò, L.; Vetri, V. Macromolecular crowding modulates phase behaviour and structural maturation of *β*-lactoglobulin condensates. International Journal of Biological Macromolecules 2026, 380, 154075.

(36) Singha, S.; Kim, D.; Roy, B.; Sambasivan, S.; Moon, H.; Rao, A. S.; Kim, J. Y.; Joo, T.; Park, J. W.; Rhee, Y. M.; others A structural remedy toward bright dipolar fluorophores in aqueous media. Chemical science 2015, 6, 4335–4342.

(37) Biancalana, M.; Makabe, K.; Koide, A.; Koide, S. Molecular mechanism of thioflavin-T binding to the surface of *β*-rich peptide self-assemblies. Journal of molecular biology 2009, 385, 1052–1063.

(38) Sancataldo, G.; Anselmo, S.; Vetri, V. Phasor-FLIM analysis of thioflavin T self-quenching in Concanavalin amyloid fibrils. Microscopy Research and Technique 2020, 83, 811–816.

(39) Anselmo, S.; Sancataldo, G.; Vetri, V. Deciphering amyloid fibril molecular maturation through FLIM-phasor analysis of thioflavin T. Biophysical Reports 2024, 4.

(40) Moreno Cerezo, F.; Corrales Rocafort, S.; Sevilla Sierra, P.; García-Blanco, F.; Díaz Oliva, C.; Catalán, J. Photophysical study of the probes acrylodan, ANS and prodan in aqueous mixtures of various alcohols. Helvetica Chimica Acta 2001, 84, 3306–3312.

(41) Rowe, B. A.; Roach, C. A.; Lin, J.; Asiago, V.; Dmitrenko, O.; Neal, S. L. Spectral heterogeneity of PRODAN fluorescence in isotropic solvents revealed by multivariate photokinetic analysis. The Journal of Physical Chemistry A 2008, 112, 13402–13412.

(42) Green, A. M.; Naughton, H. R.; Nealy, Z. B.; Pike, R. D.; Abelt, C. J. Carbonyl-twisted 6-acyl-2-dialkylaminonaphthalenes as solvent acidity sensors. The Journal of Organic Chemistry 2013, 78, 1784–1789.

(43) Yoon, A. H.; Whitworth, L. C.; Wagner, J. D.; Abelt, C. J. 2,5-PRODAN derivatives as highly sensitive sensors of low solvent acidity. Molecules 2014, 19, 6415–6427.

(44) Pantoja, C. F.; Ibáñez de Opakua, A.; Cima-Omori, M.-S.; Zweckstetter, M. Determining the physico-chemical composition of biomolecular condensates from spatially-resolved NMR. Angewandte Chemie International Edition 2023, 62, e202218078.

(45) McCall, P. M.; Kim, K.; Shevchenko, A.; Ruer-Gruß, M.; Peychl, J.; Guck, J.; Shevchenko, A.; Hyman, A. A.; Brugués, J. A label-free method for measuring the composition of multicomponent biomolecular condensates. Nature Chemistry 2025, 17, 1891–1902.

(46) Sahu, M. S. R.; Wei, J.; Weitz, D. A.; Knowles, T. P. J.; Garai, K. Liquid-solid coexistence at single fibril resolution during tau condensate ageing. bioRxiv 2025, Preprint.

(47) Zhang, W.; Falcon, B.; Murzin, A. G.; Fan, J.; Crowther, R. A.; Goedert, M.; Scheres, S. H. W. Heparin-induced tau filaments are polymorphic and differ from those in Alzheimer’s and Pick’s diseases. eLife 2019, 8, e43584.

(48) Torrado, B.; Malacrida, L.; Ranjit, S. Linear Combination Properties of the Phasor Space in Fluorescence Imaging. Sensors 2022, 22, 999.

(49) Ye, S.; Latham, A. P.; Tang, Y.; Hsiung, C.-H.; Chen, J.; Luo, F.; Liu, Y.; Zhang, B.; Zhang, X. Micropolarity governs the structural organization of biomolecular condensates. Nature Chemical Biology 2024, 20, 443–451.

(50) Ferrari, L.; Rüdiger, S. G. D. Recombinant production and purification of the human protein Tau. Protein Engineering, Design and Selection 2018, 31, 447–455.

(51) Schrimpf, W.; Barth, A.; Hendrix, J.; Lamb, D. C. PAM: A Framework for Integrated Analysis of Imaging, Single-Molecule, and Ensemble Fluorescence Data. Biophysical Journal 2018, 114, 1518–1528.

(52) Stone, M. Cross-Validatory Choice and Assessment of Statistical Predictions. Journal of the Royal Statistical Society: Series B (Methodological) 1974, 36, 111–133.

(53) Roberts, D. R.; Bahn, V.; Ciuti, S.; Boyce, M. S.; Elith, J.; Guillera-Arroita, G.; Hauenstein, S.; Lahoz-Monfort, J. J.; Schröder, B.; Thuiller, W.; Warton, D. I.; Wintle, B. A.; Hartig, F.; Dormann, C. F. Cross-validation strategies for data with temporal, spatial, hierarchical, or phylogenetic structure. Ecography 2017, 40, 913–929.

