## Supporting Information for "Microenvironmental Fingerprint of Maturing Tau Condensates from Spectral and Lifetime Phasor Analysis of a Solvatochromic Dye"

*¶imec, IUMAT, Wetenschapspark 1, B-3590 Diepenbeek, Belgium*

*§BIOMED Biomedical Research Institute, Faculty of Medicine and Life Sciences, Hasselt University, Diepenbeek, Belgium*

*‖Soft Matter and Biophysics, KU Leuven, Celestijnenlaan 200D, B-3001 Leuven, Belgium*

#### Contents

Supplementary Methods – Photophysical characterization of ACDAN in reference solvents

Supplementary Note 1 – Strickler–Berg analysis

Supplementary Note 2 – Cross-validated comparison of spectral and lifetime coordinates

Supplementary Figure S1 – Chemical structures of ACDAN and Thioflavin T

Supplementary Figure S2 – Absorption spectra in the reference solvents

Supplementary Table S1 – Individual quantum yield determinations

Supplementary Table S2 – Photophysical parameters

Supplementary Table S3 – Strickler–Berg analysis

Supplementary Table S4 – Cross-validated prediction of the spectral and lifetime coordinates

### Supplementary Methods

#### Sample preparation

ACDAN (2.5 mg) was dissolved in DMSO to prepare a 5 mM stock solution, from which 1 µM working solutions were prepared by dilution in Milli-Q water (Merck Milli-Q® Benchtop system), ethylene glycol, glycerol, and absolute ethanol. All measurements were performed on freshly prepared samples in 1-cm quartz cuvettes under ambient conditions.

#### Steady-state absorption and emission

Electronic absorption spectra were recorded on a Varian Cary 5000 UV-Vis-NIR spectrophotometer (Agilent Technologies). Corrected steady-state emission spectra were collected on an Edinburgh Instruments FLS1000 spectrofluorometer equipped with a 450 W Xenon arc lamp, using an excitation wavelength of 360 nm (maximum absorption of ACDAN in water) with 1 nm bandwidth slits for both excitation and emission, and a PMT-980 detector in right-angle geometry.

#### Time-resolved fluorescence

Fluorescence lifetime decays were recorded by time-correlated single photon counting (TCSPC) using a 375 nm picosecond pulsed diode laser (EPL series, Edinburgh Instruments; measured peak wavelength 377.6 nm) at a repetition rate of 20 MHz (pulse period 50 ns; pulse width 60 ps), with emission collected at 525 nm through a 1 nm bandwidth slit. The instrument response function (IRF) was measured using LUDOX colloidal silica as a scattering reference and used for reconvolution fitting. Decay traces were analyzed with a monoexponential model, and goodness of fit was assessed by the reduced $\chi^2$ statistic (Supplementary Table S2).

#### Absolute quantum yield

Absolute photoluminescence quantum yields ($\Phi$) were determined using the integrating sphere accessory of the FLS1000 at an excitation wavelength of 360 nm, with a cuvette of pure solvent as the reference blank. Emission was scanned from 340 to 840 nm in 1 nm steps, capturing both the scattered excitation band and the sample emission in a single scan. Quantum yields are reported as the mean of at least three independent determinations (Supplementary Table S1).[S1]

### Supplementary Note 1

#### Strickler–Berg analysis

The natural radiative lifetime $\tau_0$ was estimated from the absorption and emission spectra using the Strickler–Berg relation,[S2,S3] expressed in the wavelength domain as

$$\frac{1}{\tau_0} = 2.88 \times 10^{-9}\, n^2 \, \frac{\displaystyle\int I(\lambda)\,\mathrm{d}\lambda}{\displaystyle\int I(\lambda)\,\lambda^3\,\mathrm{d}\lambda} \times 10^{21} \int \frac{\varepsilon(\lambda)}{\lambda}\,\mathrm{d}\lambda, \tag{S1}$$

where $n$ is the solvent refractive index (water: 1.333; ethylene glycol: 1.432; glycerol: 1.475; ethanol: 1.361[S4]), $\varepsilon(\lambda)$ is the molar extinction coefficient ($M^{-1}cm^{-1}$) derived from absorbance measurements with a 1 cm path length, and $I(\lambda)$ is the normalized emission intensity. With wavelengths expressed in nm, the factor $10^{21}$ converts the emission integral ratio from $nm^{-3}$ to $cm^{-3}$. The emission integral ratios reported in Supplementary Table S3 already include this conversion. The absorption integral was evaluated over 300–600 nm (Supplementary Fig. S2) and the emission integral over 400–700 nm, after baseline correction. Negative values arising from baseline subtraction were set to zero in both cases. The Strickler–Berg-derived radiative lifetime is denoted here as $\tau_{SB}$. The lifetime predicted by combining this radiative estimate with the measured quantum yield was calculated as

$$\tau_{\text{pred}}^{(\text{SB},\Phi)} = \Phi\, \tau_{\text{SB}}. \tag{S2}$$

Agreement between $\tau_{\text{pred}}^{(\text{SB},\Phi)}$ and the measured $\tau_{\text{exp}}$ is closer in the three organic solvents, where the predicted values exceed the measured ones by 6% in ethylene glycol, 15% in ethanol and 28% in glycerol, than in water, where the predicted value exceeds the measured one by more than a factor of two (Supplementary Table S3). Water is also the solvent in which ACDAN displays the largest wavenumber shift between the absorption and emission maxima ($\Delta\bar{\nu} = 8609$ cm$^{-1}$, against 6647–7072 cm$^{-1}$ in the organic solvents) and the highest apparent non-radiative rate (Supplementary Table S2). The Strickler–Berg relation assumes a sufficiently strong allowed transition and an approximately configuration-independent electronic transition moment.[S2] Excited-state reorganization, including changes in solvent interactions, can limit this approximation. The values in Supplementary Table S3 are therefore treated as approximate spectroscopic estimates. The present measurements do not establish the origin of the larger discrepancy in water.

### Supplementary Note 2

#### Cross-validated comparison of spectral and lifetime coordinates

The joint $(\theta, \tau_P)$ representation does not assume that the two coordinates are independent. The analysis below tests how well either coordinate can be reconstructed from the other in a field of view that was not used to estimate their relationship.

Bidirectional leave-one-field-out cross-validation was performed on the exported per-pixel values of $\theta$ and $\tau_P$. At each fold, all pixels belonging to one field of view were withheld, and an ordinary least-squares linear model

$$\hat{y}_i^{(-f)} = a_{-f} + b_{-f} x_i \tag{S3}$$

was fitted using the remaining fields, where $f$ denotes the withheld field. The fitted relationship was then used to predict every pixel in the excluded field. The procedure was repeated with $x = \theta$, $y = \tau_P$ and with $x = \tau_P$, $y = \theta$.

Prediction performance was calculated from the concatenated out-of-fold predictions as

$$R^2_{\mathrm{CV}} = 1 - \frac{\sum_i (y_i - \hat{y}_i)^2}{\sum_i (y_i - \bar{y})^2}, \qquad \mathrm{RMSE}_{\mathrm{CV}} = \sqrt{\frac{1}{N} \sum_i (y_i - \hat{y}_i)^2}. \tag{S4}$$

The complete procedure was also repeated with a third-order polynomial, $y = a + bx + cx^2 + dx^3$, as a simple check for a smooth nonlinear relationship between the two coordinates.

Prediction of $\tau_P$ from $\theta$ gave $R^2_{\mathrm{CV}} = 0.688$ and $\mathrm{RMSE}_{\mathrm{CV}} = 0.209$ ns, whereas prediction of $\theta$ from $\tau_P$ gave $R^2_{\mathrm{CV}} = 0.685$ and $\mathrm{RMSE}_{\mathrm{CV}} = 0.087$ rad (Supplementary Table S4). The cubic models did not improve either prediction, giving $R^2_{\mathrm{CV}} = 0.684$ in both directions.

### Supplementary Figures

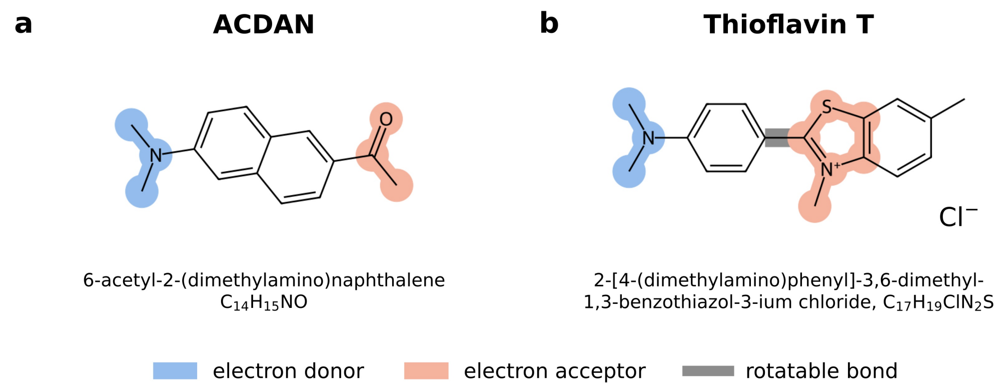


**Figure S1: Chemical structures of the fluorescent probes used in this work.** (a) ACDAN, a push–pull naphthalene derivative in which the 2-dimethylamino group acts as electron donor and the 6-acetyl group as electron acceptor. (b) Thioflavin T, shown as the benzothiazolium cation with its chloride counterion. The dimethylaminophenyl group acts as electron donor and the benzothiazolium ring as electron acceptor. The gray bond marks the central C–C bond between the two aromatic moieties, whose torsional motion provides a non-radiative decay pathway that is restricted upon binding to cross-$\beta$ structures.[S5]

### Supplementary Tables

**Table S1: Individual absolute quantum yield determinations for ACDAN.** Each entry is an independent measurement performed in the integrating sphere under identical excitation and detection settings.

| Determination | Water | Ethylene glycol | Glycerol | Ethanol |
|---|---|---|---|---|
| 1 | 0.22 | 0.69 | 0.76 | 0.83 |
| 2 | 0.21 | 0.66 | 0.78 | 0.86 |
| 3 | 0.19 | 0.65 | 0.78 | 0.83 |
| 4 | 0.18 | – | – | 0.83 |
| Mean | 0.20 | 0.67 | 0.77 | 0.84 |
| s.d. | 0.02 | 0.02 | 0.01 | 0.02 |

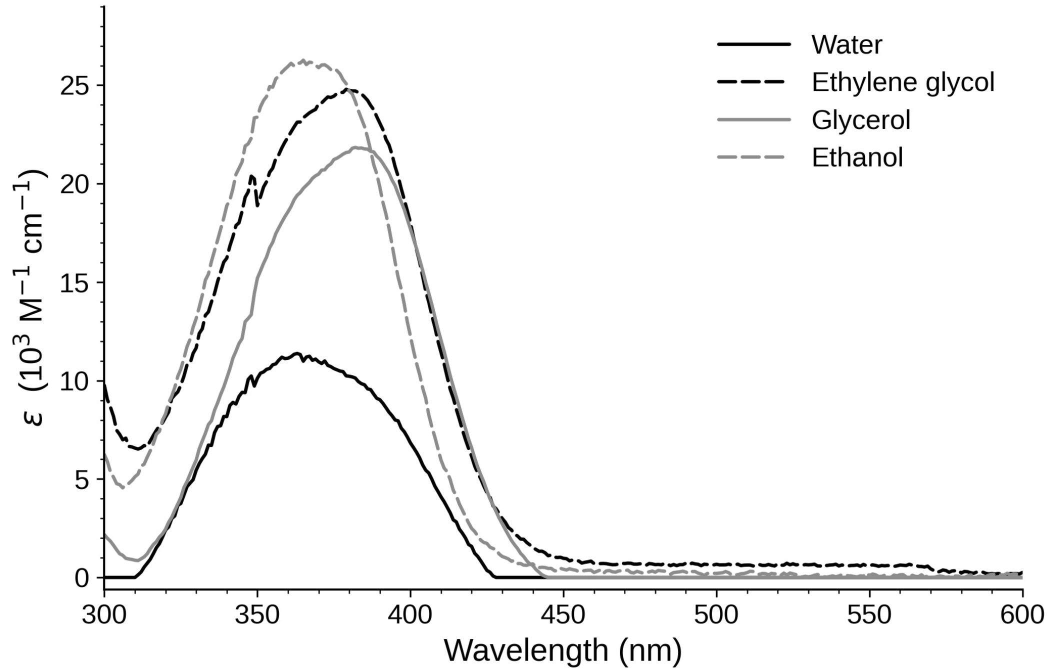


**Figure S2: Absorption spectra of ACDAN in the reference solvents.** Molar absorptivity over the 300–600 nm interval used for the absorption integral of Eq. (S1), obtained from absorbance recorded in 1 cm quartz cuvettes at the working concentrations given in the Supplementary Methods, and corrected for a baseline taken between 700 and 800 nm. Negative values arising from the baseline correction were set to zero, as in the integration. The small discontinuity near 350 nm coincides with the light-source changeover of the spectrophotometer and contributes less than 1% to the absorption integral.

**Table S2: Photophysical parameters of ACDAN in the reference solvents.** $\lambda_{\mathrm{abs}}$ and $\lambda_{\mathrm{em}}$, absorption and emission maxima; $\Delta\bar{\nu}$, wavenumber shift between the absorption and emission maxima; $n$, solvent refractive index at 20 °C; $\Phi$, absolute quantum yield (mean ± s.d.; $N$, number of independent determinations); $\tau_{\mathrm{exp}}$, lifetime obtained by monoexponential reconvolution fitting, with the associated reduced $\chi^2$; $k_r$ and $k_{nr}$, apparent radiative and non-radiative rates.

| Solvent | $\lambda_{\mathrm{abs}}$ (nm) | $\lambda_{\mathrm{em}}$ (nm) | $\Delta\bar{\nu}$ ($\mathrm{cm}^{-1}$) | $n$ | $\Phi$ ($N$) | $\tau_{\mathrm{exp}}$ (ns) | $\chi^2$ | $k_r$ / $k_{nr}$ ($\mathrm{ns}^{-1}$) |
|---|---|---|---|---|---|---|---|---|
| Water | 363 | 528 | 8609 | 1.333 | $0.20 \pm 0.02$ (4) | 1.24 | 5.3 | 0.16 / 0.64 |
| Ethylene glycol | 379 | 510 | 6777 | 1.432 | $0.67 \pm 0.02$ (3) | 2.75 | 1.7 | 0.24 / 0.12 |
| Glycerol | 382 | 512 | 6647 | 1.475 | $0.77 \pm 0.01$ (3) | 3.26 | 1.6 | 0.24 / 0.069 |
| Ethanol | 365 | 492 | 7072 | 1.361 | $0.84 \pm 0.02$ (4) | 3.36 | 1.3 | 0.25 / 0.048 |

**Table S3: Strickler–Berg analysis of ACDAN in the reference solvents.** $\varepsilon_{\max}$, molar extinction coefficient at the absorption maximum; $\tau_{\mathrm{SB}}$, radiative lifetime from Eq. (S1); $\tau_{\mathrm{pred}}^{(\mathrm{SB},\Phi)} = \Phi\,\tau_{\mathrm{SB}}$; $\Delta$, deviation of the predicted from the measured lifetime of Supplementary Table S2. Absorbance was converted to molar absorptivity using the working concentrations given in the Supplementary Methods. Molar absorptivities were obtained from a single absorbance measurement per solvent and are therefore rounded to the nearest 100 $\mathrm{M^{-1}cm^{-1}}$.

| Solvent | $\varepsilon_{\max}$ ($\mathrm{M^{-1}cm^{-1}}$) | $\int \varepsilon\,\mathrm{d}\lambda/\lambda$ ($\mathrm{M^{-1}cm^{-1}}$) | $\int I\,\mathrm{d}\lambda/\int I\lambda^3\mathrm{d}\lambda$ ($\mathrm{cm^{-3}}$) | $\tau_{\mathrm{SB}}$ (ns) | $\tau_{\mathrm{pred}}^{(\mathrm{SB},\Phi)}$ (ns) | $\tau_{\exp}$ (ns) | $\Delta$ (%) |
|---|---|---|---|---|---|---|---|
| Water | 11,400 | $2.19 \times 10^3$ | $6.22 \times 10^{12}$ | 14.35 | 2.87 | 1.24 | +131 |
| Ethylene glycol | 24,800 | $5.69 \times 10^3$ | $6.80 \times 10^{12}$ | 4.38 | 2.92 | 2.75 | +6 |
| Glycerol | 21,900 | $4.22 \times 10^3$ | $7.01 \times 10^{12}$ | 5.40 | 4.18 | 3.26 | +28 |
| Ethanol | 26,300 | $5.34 \times 10^3$ | $7.63 \times 10^{12}$ | 4.60 | 3.86 | 3.36 | +15 |

**Table S4: Bidirectional cross-validated prediction of the spectral and lifetime coordinates.** $R^2_{\mathrm{CV}}$ and $\mathrm{RMSE}_{\mathrm{CV}}$ were calculated from the concatenated leave-one-field-out predictions. In each fold, all pixels from one field were withheld together and the model was fitted on the remaining fields.

| Predicted coordinate | Predictor/model | $R^2_{\mathrm{CV}}$ | $\mathrm{RMSE}_{\mathrm{CV}}$ |
|---|---|---|---|
| $\tau_P$ | $\theta$, linear | 0.688 | 0.209 ns |
| $\tau_P$ | $\theta$, cubic | 0.684 | 0.210 ns |
| $\theta$ | $\tau_P$, linear | 0.685 | 0.087 rad |
| $\theta$ | $\tau_P$, cubic | 0.684 | 0.088 rad |

### References

(S1) Würth, C.; Grabolle, M.; Pauli, J.; Spieles, M.; Resch-Genger, U. Relative and Absolute Determination of Fluorescence Quantum Yields of Transparent Samples. *Nat. Protoc.* **2013**, *8*, 1535–1550.

(S2) Strickler, S. J.; Berg, R. A. Relationship between Absorption Intensity and Fluorescence Lifetime of Molecules. *J. Chem. Phys.* **1962**, *37*, 814–822.

(S3) Lakowicz, J. R. *Principles of Fluorescence Spectroscopy*, 3rd ed.; Springer: New York, 2006.

(S4) Lide, D. R., Ed. *CRC Handbook of Chemistry and Physics*, 90th ed.; CRC Press: Boca Raton, 2009.

(S5) Biancalana, M.; Makabe, K.; Koide, A.; Koide, S. Molecular Mechanism of Thioflavin-T Binding to the Surface of $\beta$-Rich Peptide Self-Assemblies. *J. Mol. Biol.* **2009**, *385*, 1052–1063.
